# Polygenicity of complex traits is explained by negative selection

**DOI:** 10.1101/420497

**Authors:** Luke J. O’Connor, Armin P. Schoech, Farhad Hormozdiari, Steven Gazal, Nick Patterson, Alkes L. Price

## Abstract

Complex traits and common disease are highly polygenic: thousands of common variants are causal, and their effect sizes are almost always small. Polygenicity could be explained by negative selection, which constrains common-variant effect sizes and may reshape their distribution across the genome. We refer to this phenomenon as *flattening*, as genetic signal is flattened relative to the underlying biology. We introduce a mathematical definition of polygenicity, the *effective number of associated SNPs*, and a robust statistical method to estimate it. This definition of polygenicity differs from the number of causal SNPs, a standard definition; it depends strongly on SNPs with large effects. In analyses of 33 complex traits (average N=361k), we determined that common variants are ∼4x more polygenic than low-frequency variants, consistent with pervasive flattening. Moreover, functionally important regions of the genome have increased polygenicity in proportion to their increased heritability, implying that heritability enrichment reflects differences in the number of associations rather than their magnitude (which is constrained by selection). We conclude that negative selection constrains the genetic signal of biologically important regions and genes, reshaping genetic architecture.

## Introduction

Genome-wide association studies (GWAS) have revealed that common diseases and complex traits are heritable and highly polygenic^1-12^. There are usually no large-effect common SNPs, and hundreds or thousands of small-effect SNPs are required to explain a large proportion of heritability. This aspect of polygenicity presents a challenge for geneticists, as small-effect SNPs are difficult to detect (leading to “missing heritability”^13^) and difficult to interpret^14,15^. The large number of causal SNPs could be explained by extraordinary biological complexity, e.g. if thousands of genes affect a trait under an “omnigenic model”^15^. However, biological complexity does not explain the absence of large-effect SNPs.

Here, we investigate a complementary hypothesis: due to negative selection, large-effect SNPs are prevented from becoming common in the population while small-effect SNPs are unaffected, resulting in increased polygenicity for common variants (Figure 1). We refer to this phenomenon as *flattening*, as the distribution of heritability across the genome is flattened relative to the underlying biology. Although it is widely known that negative selection constrains common-variant effect sizes on average^10,16-20^, the extent to which negative selection leads to increased polygenicity is unknown.

**Figure 1.**
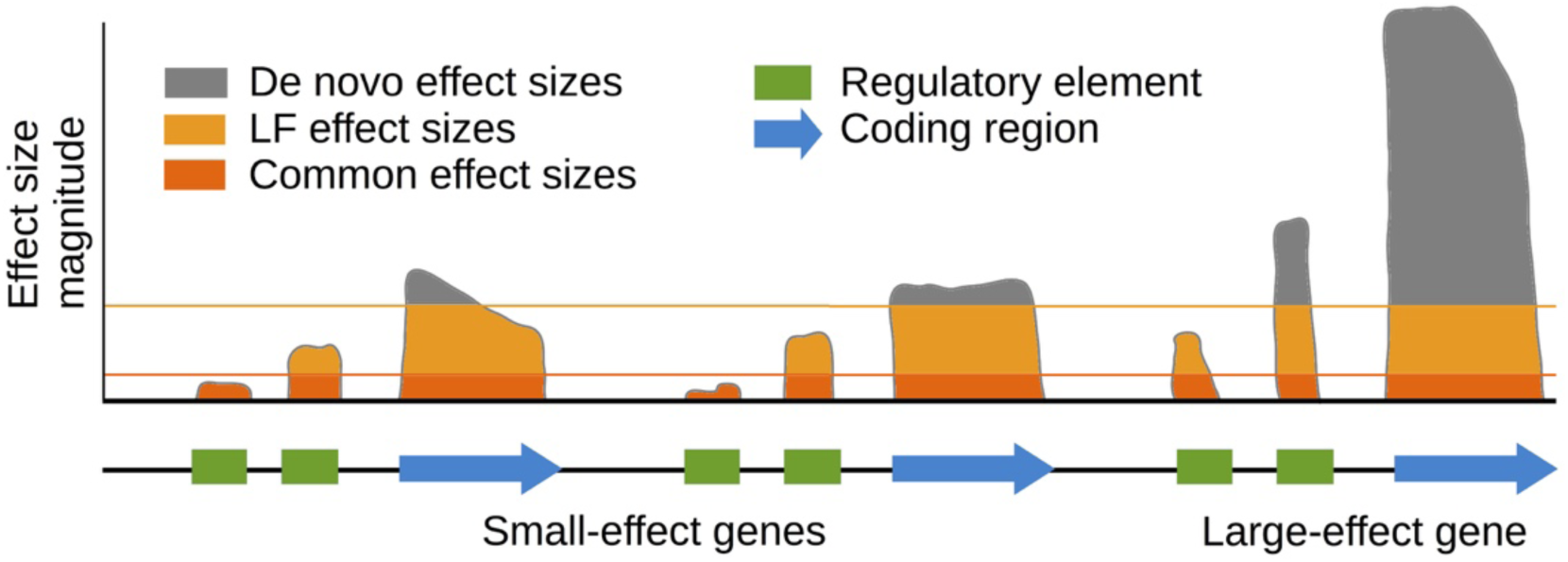
Illustration of flattening due to natural selection. We display the maximum per-allele effect size of a SNP at each site for a toy example of three genes and nearby regulatory regions. Here, the distribution of *de novo* effects is not highly polygenic; it is dominated by coding variants in a single large-effect gene (although other genes also harbor small effects). Negative selection imposes an upper effect size bound (possibly soft), resulting in increased polygenicity. The effect is stronger for common SNPs than for low-frequency SNPs. Within functionally important regions (e.g. coding), a larger proportion of variants have effect sizes near the bound, leading to especially large polygenicity. In practice, this bound may vary across the genome; we expect that the resulting distribution will be more flat than the distribution of *de novo* variants, but not perfectly flat.

Flattening would lead to increased polygenicity for common vs. low-frequency SNPs and especially high polygenicity within functionally important regions of the genome (Figure 1). In order to compare polygenicity across allele frequencies and functional categories, we introduced a mathematical definition of polygenicity and developed a method, stratified LD fourth moments regression (S-LD4M), to estimate it using summary association statistics. We applied S-LD4M to summary statistics for 33 diseases and complex traits (average N=361k).

## Overview of methods

In order to quantify the effects of flattening, we introduce a mathematical definition of polygenicity, the *effective number of associated SNPs* (*M*_*a*_). This quantity describes the number of genetic associations that explain the heritability of a trait. It differs from the total number of causal SNPs (*M*_*c*_): if a small number of SNPs explain a large proportion of heritability, and a large number of SNPs have extremely small effect sizes, then *M*_*a*_ will be much smaller than *M*_*c*_.

If there is no linkage disequilibrium (LD) between causal SNPs, then *M*_*a*_ is inversely proportional to the normalized fourth moment, or kurtosis *κ*, of the causal effect size distribution:

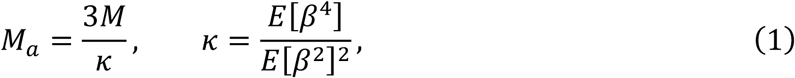

where *M* is the number of SNPs and *β* is the causal effect size of a SNP in per-normalized-genotype units (i.e. per-allele effect size times 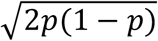, where *p* is the allele frequency). SNPs with large effects contribute strongly to E[*β*^4^], leading to decreased *M*_*a*_. In the presence of LD between causal SNPs, *M*_*a*_ is defined using mixed fourth moments of causal and marginal effect sizes, ensuring that *M*_*a*_ is identifiable (Methods). Under an infinitesimal (Gaussian) architecture, *M*_*a*_ is equal to the effective number of independent SNPs^21^.

We also define the *average unit of heritability*, denoted 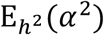, as the marginal per-SNP heritability averaged across components of heritability (not uniformly across SNPs). 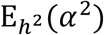 multiplied by *M*_*a*_ is equal to h^2^. For example, in the special case that causal SNPs are independent and their effect sizes follow a *N*(0, *σ*^2^) distribution, 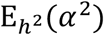 is equal to *σ*^2^ and *M*_*a*_ is equal to the number of causal SNPs. More generally, 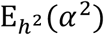 can be visualized as the area of the shaded region in Figure 2a, and it depends strongly on large-effect SNPs (Figure 2b). For example, if 4 causal SNPs each contribute 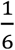 of heritability (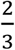 of heritability in total) and 96 other causal SNPs have much smaller effects, then the average unit of heritability is approximately 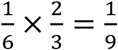, and *M*_*a*_ ≈ 9 (Figure 2a bottom).

**Figure 2.**
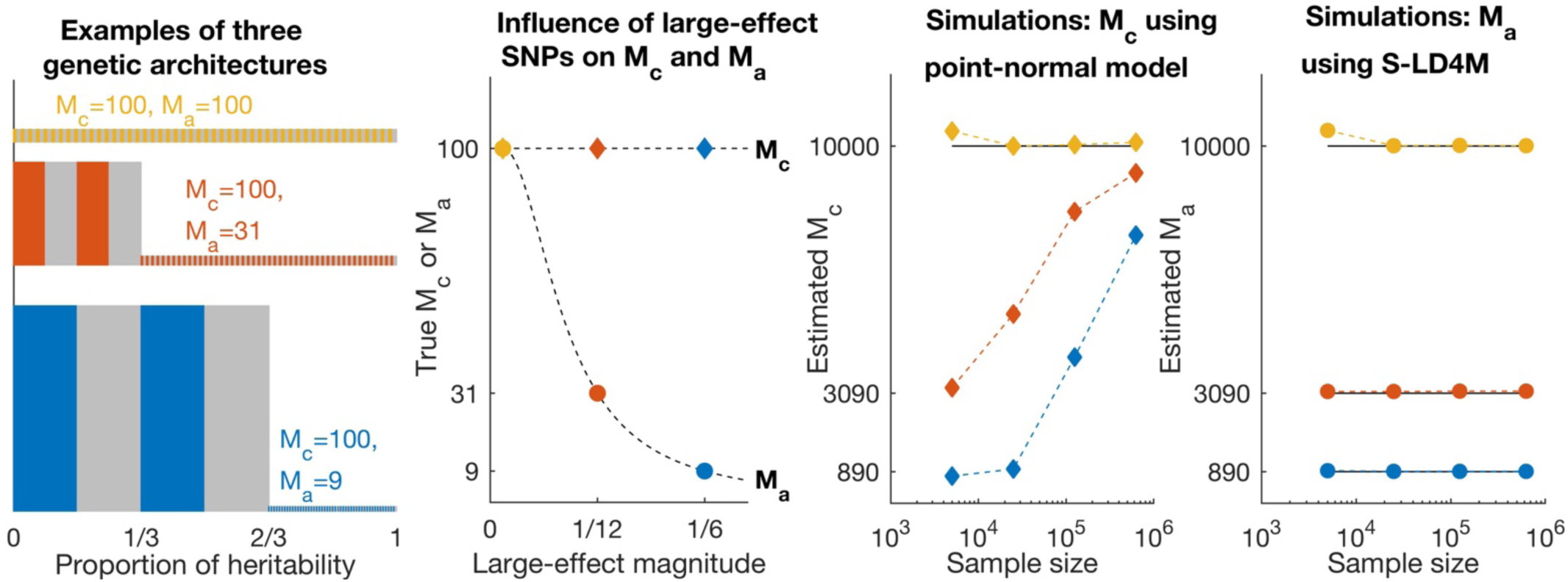
Comparison of the effective number of associated SNPs (*M*_*a*_) with the number of causal SNPs (*M*_*c*_). (a),(b) Examples of three genetic architectures with *M*_*c*_ = 100. (a) Each colored or gray block corresponds to one SNP; both height and width are proportional to the SNP’s effect size variance. The average unit of heritability, denoted 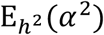, is the average height (equal to the total area) of the colored and gray regions. *M*_a_ is equal to 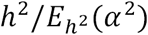 (b)*M*_*c*_ and *M*_a_ as a function of the effect size magnitude of the 4 large-effect SNPs. (c),(d) Simulations of the same three genetic architectures with the number of SNPs (and causal SNPs) scaled up by 100x. (c) Estimates of *M*_*c*_ under a point-normal model, at different sample sizes. (d) Estimates of *M*_a_ using S-LD4M, at different sample sizes. Error bars denote 95% confidence intervals (based on 1,000 simulations), but are smaller than the data points. Numerical results are reported in Supplementary Table 1.

An alternative definition of polygenicity, which is widely used, is the total number of causal SNPs (*M*_*c*_)^1,3,5-7,10-12^. We chose to estimate *M*_*a*_ rather than *M*_*c*_ for three reasons. First, *M*_*c*_ is difficult to estimate robustly, as it is impossible to distinguish SNPs with zero effect from SNPs with arbitrarily small effect (see Simulations). Second, because negative selection affects large-effect SNPs more strongly than small-effect SNPs, it is expected to influence *M*_*a*_ much more strongly than *M*_*c*_ (Figure 2b). Third, *M*_*a*_ (but not *M*_*c*_) is closely related to missing heritability and polygenic prediction accuracy (Methods).

**Table 1.**
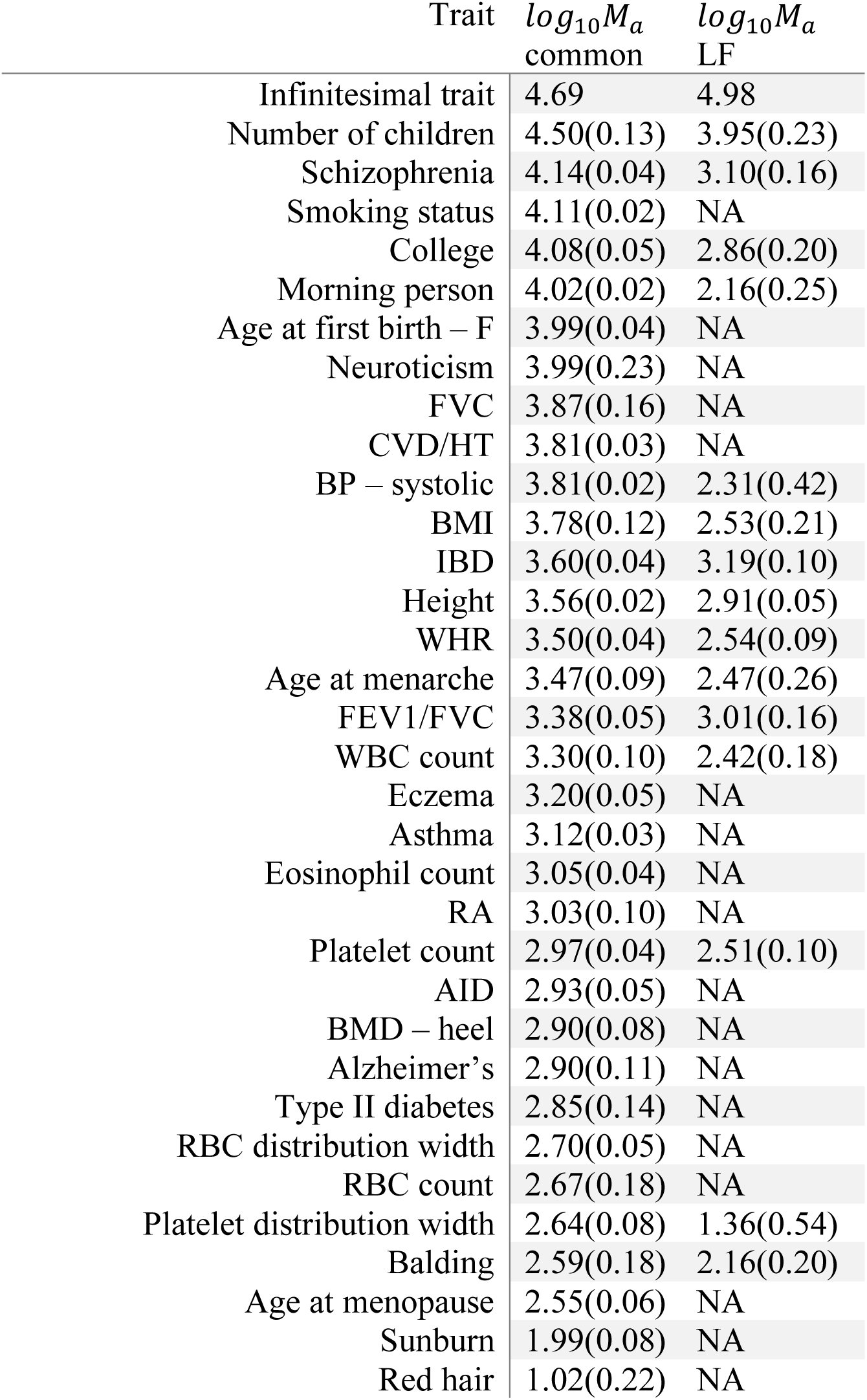
Estimates of polygenicity for common and low-frequency SNPs across 33 complex traits. We report common variant estimates for all traits, and low-frequency estimates for well-powered traits (see Methods). The first row reports the effective number of independent SNPs^21^. *M*_a_ is close to this value when marginal effect sizes approximately follow a Normal distribution, which does not imply that every SNP is causal.

*M*_*a*_ can be defined for categories of SNPs, such as low-frequency SNPs or coding SNPs. To compare categories of different size, we divide *M*_*a*_ by the number of SNPs in each category. We refer to differences in per-SNP *M*_*a*_ simply as differences in polygenicity. We define *polygenicity enrichment* as the per-SNP *M*_*a*_ of all SNPs in a category divided by the per-SNP *M*_*a*_ of all SNPs; analogously, we define *heritability enrichment* as the per-SNP heritability of all SNPs in a category divided by the per-SNP heritability of all SNPs (similar to previous work^22^). “All SNPs” refers to common (*MAF* ≥ 0.05) and low-frequency (0.005 ≤ *MAF <* 0.05) SNPs. Enrichment can be either >1 or <1 (i.e. depletion).

We developed a method, stratified LD fourth moments regression (S-LD4M), to estimate *M*_*a*_ from summary association statistics and LD from a reference panel. S-LD4M regresses squared χ^2^ statistics (i.e. fourth powers of signed Z scores) on *LD fourth moments*, defined as sums of 4 values to each category of SNPs. This approach is analogous to stratified LD score regression (S-LDSC), which regresses χ^2^ statistics on LD scores (LD second moments)^22^. We use a slightly modified version of S-LDSC to estimate heritability enrichment. We run S-LD4M and S-LDSC using the baseline-LD model^17^, which includes 75 coding, conserved, regulatory, MAF- and LD-related annotations. When estimating the polygenicity and polygenicity enrichment of a category of SNPs, we restrict to well-powered traits. Further details are provided in the Methods section. We have released open-source software implementing S-LD4M (see URLs).

## Simulations

First, we performed simple simulations with no LD to evaluate both the S-LD4M estimator of *M*_*a*_ (the effective number of associated SNPs), and a maximum-likelihood estimator of *M*_*c*_ (the number of causal SNPs) under a point-normal model (PN) (see Methods). This estimator is expected to produce similar results as previous estimators of *M*_*c*_ under a point-normal model^3,7,9-12^. We simulated 400 large-effect SNPs and 9,600 small-effect SNPs, with *M* = 50k total SNPs. The mixture of different causal effect sizes, violating the point-normal model, is consistent with evidence for real traits^10,11^. When there was a large difference between the causal effect sizes of large- and small-effect SNPs, PN produced biased and sample size-dependent estimates of *M*_*c*_ (Figure 2c and Supplementary Table 1), consistent with recent work^11^. In general, any method that estimates *M*_*c*_ will only detect causal SNPs with effect sizes greater than some threshold, where that threshold depends on modeling assumptions and power. In contrast, S-LD4M does not depend on parametric modeling assumptions, and it produces unbiased estimates of *M*_*a*_ that do not depend on power (Figure 2d). Power-dependent bias would be especially problematic for comparing common vs. low frequency polygenicity, due to lower power for low-frequency SNPs.

Next, we performed a series of simulations using real LD patterns to determine whether S-LD4M produces reliable estimates of polygenicity in realistic settings. We simulated summary association statistics from the asymptotic sampling distribution^23^ for UK Biobank imputed SNPs on chromosome 1 (*M ≈* 1.0M SNPs). We used *N* = 50k samples and 2= 0.2, to approximately match 2*/M* (and hence the expected χ^2^ statistic) for UK Biobank traits. We included MAF-dependent genetic architectures^10,20^ and causal effect size heterogeneity. Details of each simulation are provided in the Methods section.

First, we assessed the ability of S-LD4M to estimate polygenicity for the set of all common and low-frequency SNPs. We determined that median estimates of *M*_*a*_ were approximately unbiased across a wide range of true values, although slight bias was observed for very low values of *M*_*a*_ (Figure 3a and Supplementary Table 1). We report medians instead of means due to noise in the denominator of the estimator, which leads to instability in the mean. In general, S-LD4M is expected to produce approximately unbiased median estimates given sufficient power; the bias that we observed at low values of *M*_*a*_ results from lower power when polygenicity is low (see below for simulations at higher or lower power).

**Figure 3.**
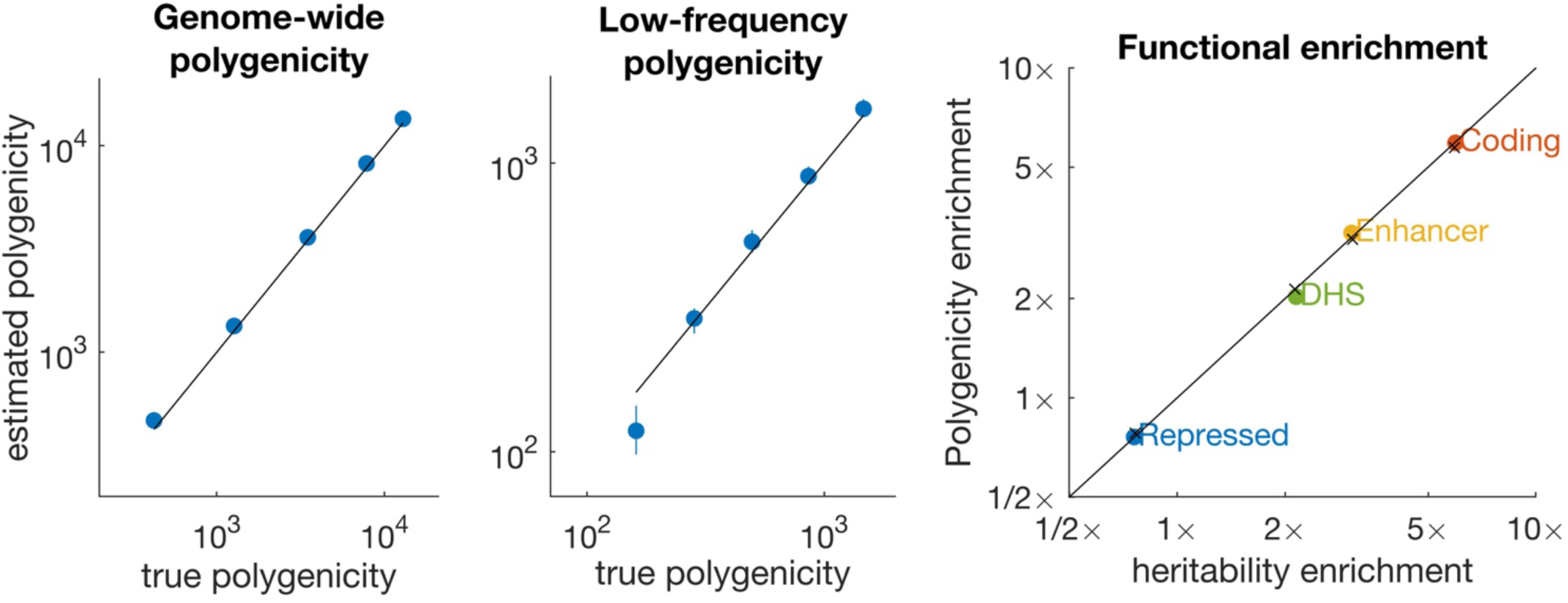
Accuracy of S-LD4M estimates in simulations with LD. (a) Estimates of *M*_a_ for all SNPs (*MAF* = 0.5 − 50%). (b) Estimates of *M*_a_ for low-frequency SNPs (*MAF* = 0.5 *−* 5%); common-SNP *M*_a_ is fixed at ∼1,000 in these simulations. (c) Estimates of polygenicity enrichment and heritability enrichment in simulations with four functional categories. Black lines denotes y=x, and colored points denote estimates. In panel (c), × denotes true values. Error bars denote 95% confidence intervals (based on 1,000 simulations), but are smaller than the data points in most cases. Numerical results are reported in Supplementary Table 1.

Second, we fixed the polygenicity of common SNPs (MAF > 5%) at *M*_*a*_ ≈ 1,000 and progressively reduced the polygenicity of low-frequency SNP (MAF = 0.5 *−* 5%). Similar to Figure 3a, median estimates of low-frequency *M*_*a*_ were approximately unbiased except for very low values of *M*_*a*_ (Figure 3b). These estimates indicate that S-LD4M can be used to compare the polygenicity of common and low-frequency SNPs.

Third, we simulated equal heritability enrichment and polygenicity enrichment in four functional categories: coding (∼8x enriched), enhancer (∼5x), DNase I hypersensitivity sites (DHS, ∼2x), and repressed (∼0.75x), similar to our results on real traits (see below). We determined that S-LD4M produces approximately unbiased estimates of polygenicity enrichment (Figure 3c). We also considered an alternative model of heritability enrichment, under which polygenicity was approximately constant across functional categories. Estimates were approximately unbiased for enriched functional categories but biased for the repressed annotation (Supplementary Figure 1). Despite the fact that this genetic architecture is less realistic (see below), we avoid reporting estimates of polygenicity for depleted functional annotations on real traits.

Fourth, we performed simulations to assess whether genetic architectures with non-random clustering of causal SNPs would bias our estimates of polygenicity enrichment for SNPs in functional categories. Such clustering is expected in real data (e.g. due to biologically important genes), and it could potentially lead to bias, as S-LD4M uses an LD approximation whose accuracy depends on LD between causal SNPs (however, we do not assume that linked SNPs have independent causal effect sizes; see Methods). We simulated clusters of either 5 or 50 causal SNPs (on average) across three genomic length scales (10, 100, 1000 SNPs; roughly 3kb, 30kb, 300kb on average). We determined that median enrichment estimates were approximately unbiased in each case, indicating that our LD approximation is robust to non-random clustering of causal SNPs (Supplementary Figure 2).

Finally, we performed simulations at different sample sizes and included a filtering step (Methods) to exclude trait-annotation pairs with inadequate power. We performed simulations at *N*=10k, *N*=50k and *N*=250k. For each SNP set (four functional categories + low-frequency SNPs), polygenicity enrichment estimates are reported in Supplementary Figure 3, and proportions of traits retained after the filtering step for each category are reported in Supplementary Table 2. At the default setting of *N*=50k, 61% of simulated traits were retained on average across categories, and polygenicity enrichment estimates were approximately unbiased. Similar results were obtained at *N*=250k. At *N*=10k, only 8% of simulated traits were retained on average (including ∼0% of simulated traits for coding SNPs and for low-frequency SNPs), and polygenicity enrichment estimates were downward biased. Our analyses of real traits more closely correspond to *N*=50k both in terms of average χ^2^ statistic and in terms of the proportion of traits retained (49% on average across categories). Nonetheless, we avoid reporting estimates of polygenicity enrichment for annotations having well-powered enrichment estimates for fewer than 10 out of the 33 traits that we analyzed. We determined that our confidence intervals were approximately well-calibrated or conservative in these simulations (Supplementary Table 2).

## Polygenicity of common and low-frequency SNPs across 33 complex traits

We applied S-LD4M to publicly available summary association statistics for 33 diseases and complex traits (average *N* = 361k; Supplementary Table 3), including 29 UK Biobank traits^24-26^ (see URLs) and 4 additional common diseases. LD scores and LD fourth moments were computed using LD estimated from UK10K^27^ (*M* = 8.5 million SNPs after QC, MAF > 0.5%). As in previous work^17,22,28^, we excluded the major histocompatibility complex, which has unusually large effect sizes and long-range LD due to balancing selection. (This choice increases *M*_*a*_ estimates for immune-related traits.) Details of our analyses are provided in the Methods section.

For most traits, *M*_*a*_ for common SNPs (MAF > 5%) ranged between 500 and 20,000 (10^2.7^ and 10^4.3^; Table 1). Fecundity- and brain-related traits were remarkably polygenic, with *M*_*a*_ estimates greater than 10,000. For Number of children, the most polygenic trait, *M*_*a*_ was almost as large as effective number of independent SNPs, corresponding to an infinitesimal trait (log_10_ *M*_*a*_ =4.52 (0.13) vs. log_10_ *M*_e_ = 4.69). This estimate implies that most common SNPs are associated with this trait (though a much smaller fraction may be causal). Schizophrenia was also extremely polygenic (log_10_ *M*_*a*_ = 4.14 (0.04)), consistent with previous work^1,4^. Particularly high polygenicity for these traits could result from particularly strong negative selection (see below), although it is also possible that they have greater biological complexity. Red hair pigmentation and sunburn were the least polygenic traits, consistent with known large-effect common SNPs for pigmentation traits^29,30^ (however, we emphasize that the total number of causal SNPs may be much larger than *M*_*a*_ for these traits).

We compared per-SNP *M*_*a*_ estimates for common and low-frequency SNPs across 15 well-powered traits (Figure 4a and Table 1). Polygenicity was 3.9x (95%CI: 2.9 *−* 5.2x) smaller for low-frequency SNPs than for common SNPs on average, with substantial variation across traits. This difference was similar to the 3.9x (95%CI: 3.5 − 4.4x) smaller per-SNP heritability of low-frequency vs. common SNPs on average (consistent with previous estimates^10,20^); this concordance was consistent across traits (Figure 4b). This concordance suggests that the upper bound on effect size imposed by negative selection is approximately constant in units of per-SNP heritability, consistent with evolutionary modeling (see below).

**Figure 4.**
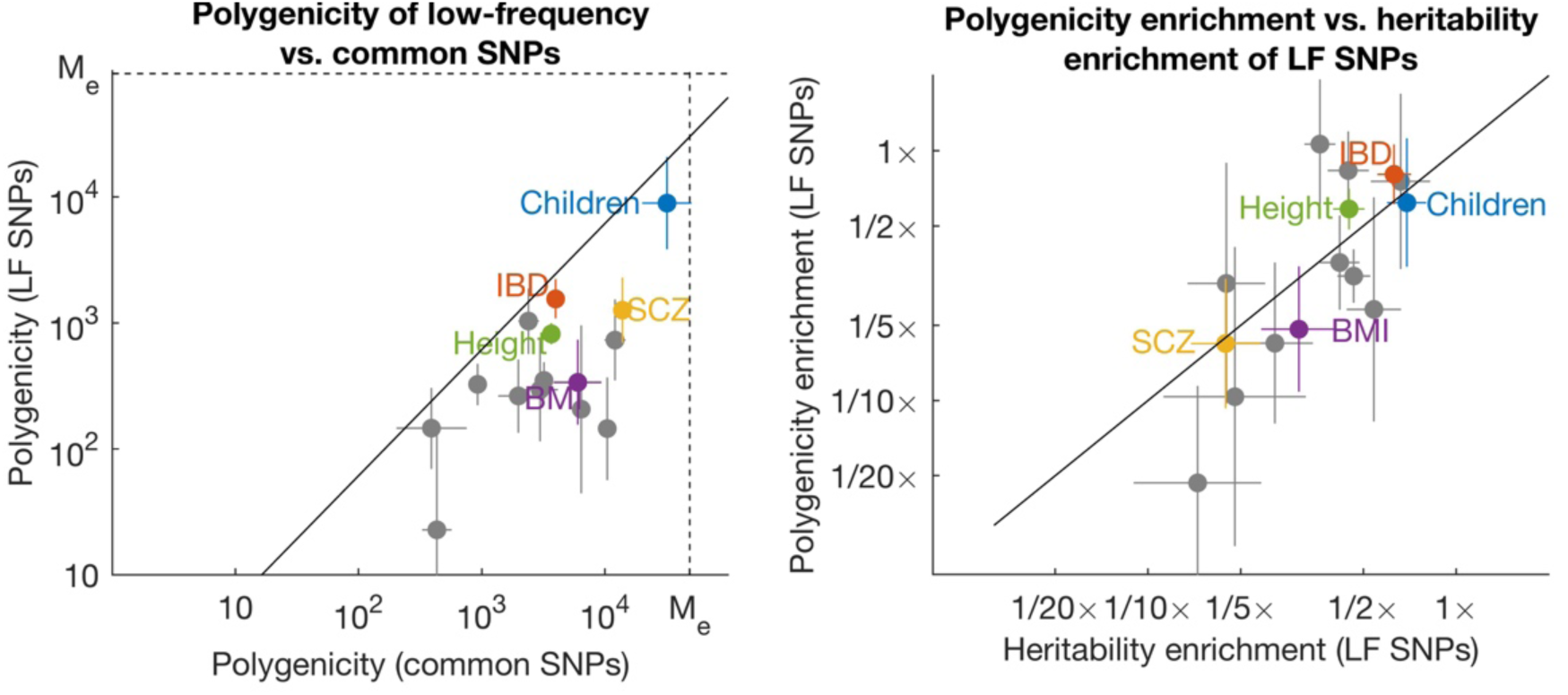
Comparison of common and low-frequency polygenicity across 15 complex traits. (a) Estimates of *M*_a_ for common and low-frequency SNPs. Estimates are meta-analyzed across well-powered traits. Common-variant polygenicity was ∼4x greater on average than low-frequency polygenicity. Dotted lines denote the effective number of independent SNPs (*M*_e_) for common and low-frequency SNPs respectively, corresponding to an infinitesimal (Gaussian) architecture. The solid line denotes equal per-SNP *M*_a_. (b) Estimates of polygenicity enrichment and heritability enrichment for low-frequency SNPs (compared to all common and low-frequency SNPs). The solid line denotes equal enrichment. Error bars denote 95% confidence intervals. Numerical results are reported in Table 1 and Supplementary Table 5.

For 6 of the 33 traits, summary association statistics from independent cohorts were available. S-LD4M produced concordant *M*_*a*_ estimates for common SNPs on these data sets, despite the smaller sample size and much smaller number of regression SNPs (Supplementary Table 4). (We did not estimate low-frequency *M*_*a*_ for these data sets because summary statistics were not available for low-frequency SNPs.) *M*_*a*_ provides an upper bound on the proportion of heritability explained by genome-wide significant loci for a trait (Methods). We compared this bound with a direct estimate of this proportion (which may be upwardly biased by winner’s curse), and determined that the predicted bound corresponded fairly closely with the estimate (Spearman *r*^2 =^ 0.61; Supplementary Figure 4a). It also provided a conservative upper bound on the number of genome-wide significant SNPs (Supplementary Figure 4b).

## Polygenicity of functional categories across 33 complex traits

We compared estimates of polygenicity enrichment with estimates of heritability enrichment across 25 main functional categories from the baseline-LD model^17^ and meta-analyzed results across well-powered traits for 21 categories with at least 10 well-powered traits (Figure 5, Supplementary Table 5 and Supplementary Table 6; 49% of trait-annotation pairs were well powered). For most annotations, polygenicity enrichment was approximately equal to heritability enrichment (regression slope = 0.93; r^2^= 0.88). For example, SNPs in conserved regions were 13x enriched for heritability and 14x enriched for polygenicity, and coding SNPs were 9.4x enriched for heritability and 6.6x enriched for polygenicity. (These functional enrichments for the union of common and low-frequency SNPs are larger than the corresponding enrichments for common SNPs, due to larger functional enrichment for low-frequency SNPs^19^.) Thus, heritability enrichment in functional categories is predominantly driven by differences in polygenicity, rather than differences in effect-size magnitude. In contrast, *de novo* SNP effect sizes are expected to be much larger in functionally important regions. Thus, genetic signals of important functional regions are constrained by negative selection (Figure 1).

**Figure 5.**
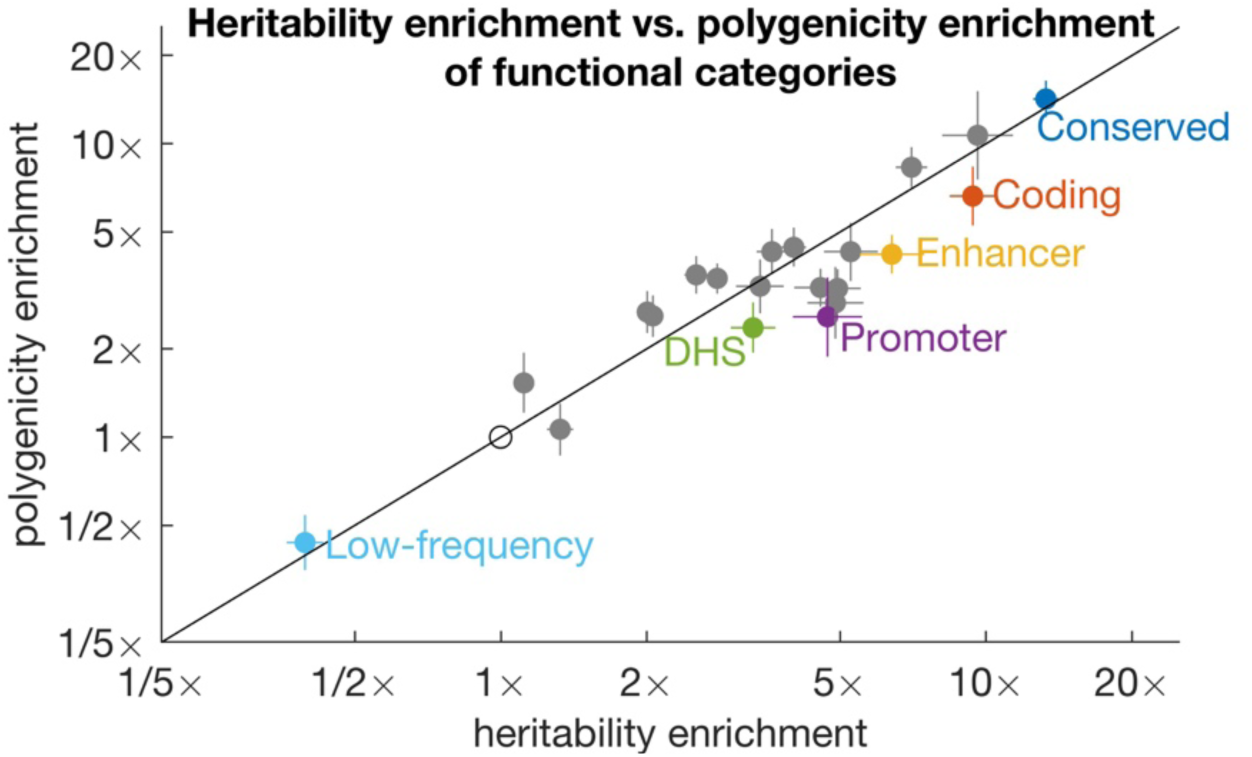
Estimates of polygenicity enrichment and heritability enrichment of functional categories. We report estimates for 20 functional categories plus low-frequency SNPs. Estimates are meta-analyzed across well-powered traits. Error bars denote 95% confidence intervals. Complete results for each trait are reported in Supplementary Table 5, and meta-analyzed results are reported in Supplementary Table 6.

We compared functional enrichment between groups of related traits (8 brain-related, 6 blood-related, and 5 immune-related traits; Supplementary Figure 5). Brain-related traits had smaller functional enrichments both for heritability and for polygenicity, consistent with previous findings^17,19,22^. Smaller functional enrichment could be explained by stronger negative selection for these traits^19^, which may strongly limit the enrichment of any functional category. Stronger negative selection would also be consistent with greater genome-wide polygenicity for these traits (Table 1).

To investigate the relationship between functional enrichment and genome-wide polygenicity, we quantified the sparsity explained by functional annotations, where sparsity is inversely proportional to *M*_*a*_ (Methods). As a proportion of total sparsity, the sparsity explained by functional annotations from the baseline-LD model ranged between 7% and 42% (Supplementary Figure 6a). Traits with smaller *M*_*a*_ (greater sparsity) had greater sparsity explained (*r*^2 =^ 0.78; Supplementary Figure 6b).

**Figure 6.**
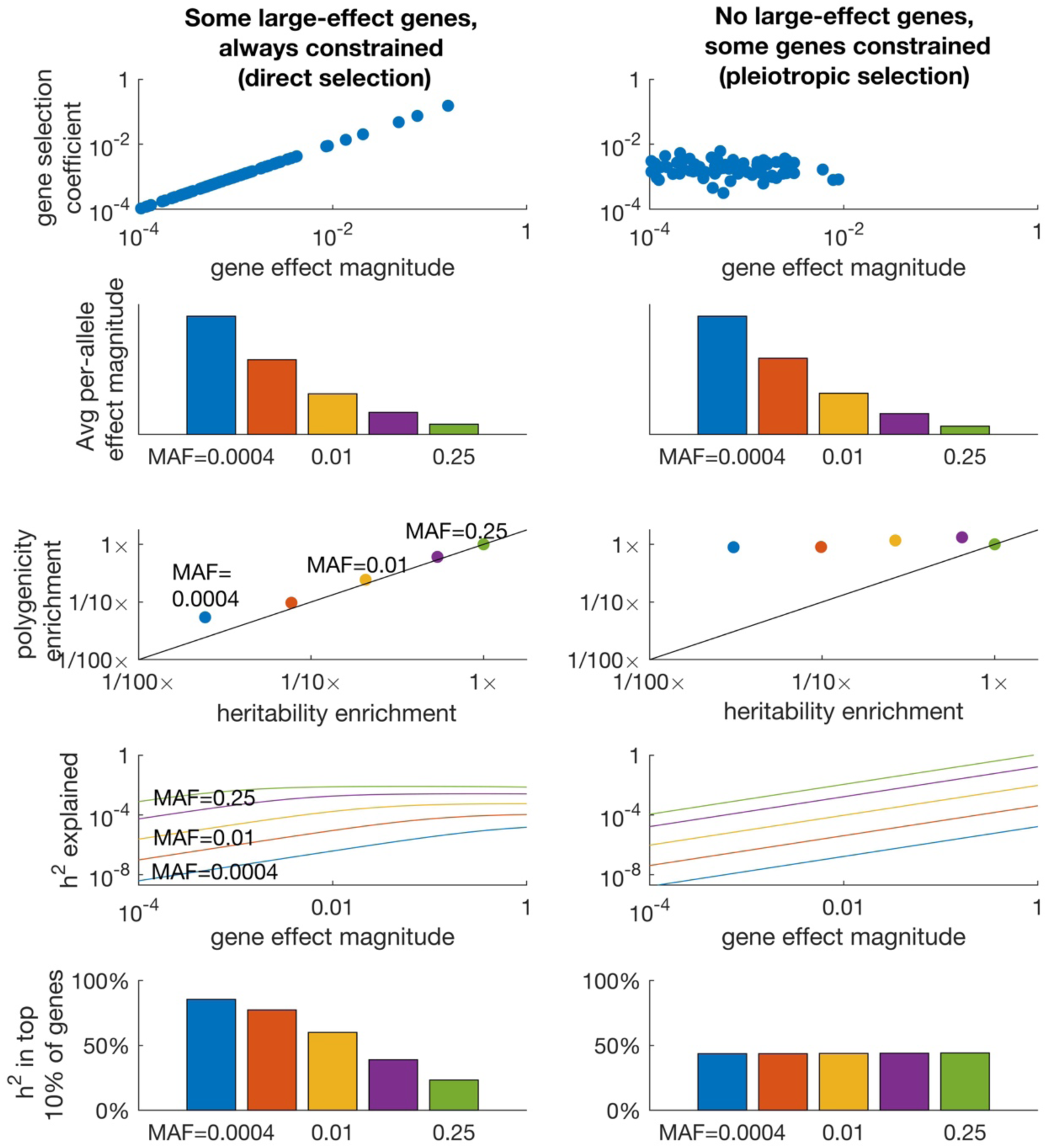
Gene-level flattening under an evolutionary model. In the left column (panels a,c,e,g,i), there are some large-effect genes, but direct stabilizing selection acting on the phenotype strongly constrains these genes. In the right column (panels b,d,f,h,j), there are no large-effect genes; pleiotropic stabilizing selection has varying effects on each gene, limiting common-SNP effect sizes on average. (a-b) Joint distribution of gene effect size magnitudes and selection coefficients. (c-d) Average squared per-allele effect sizes at different allele frequencies. The strength of selection was chosen to produce similar common-variant effect sizes in both columns. (e-f) Heritability and polygenicity enrichment at different allele frequencies (relative to MAF=0.25). Polygenicity at MAF=0.25 is approximately equal for the two columns, due to the different distributions of gene effect sizes. (g-h) Expected heritability explained by a gene as a function of its effect size for SNPs at different frequencies. (i-j) Proportion of heritability explained by the top 10% of largest-effect genes for SNPs at different allele frequencies. Numerical results are reported in Supplementary Table 7.

### GWAS signals of biologically important genes are constrained by negative selection

Flattening may lead to increased polygenicity not only at the level of SNPs, but also at the level of genes, particularly for genes with large effects on a trait. As a result, GWAS effect sizes may be similar for SNPs near small-effect genes and for SNPs near large-effect genes, and top GWAS SNPs may often implicate small-effect genes (Figure 1).

In order to investigate the impact of flattening on the distribution of heritability across genes, we explored two evolutionary fitness models: a realistic model with SNP-level flattening, and a model without SNP-level flattening that is realistic in other respects. Under both models, SNPs affect genes and genes affect the trait. Each gene has a trait effect size and a selection coefficient. In the first model (Figure 6a and Supplementary Table 7), there are large-effect genes, and these genes are always strongly constrained. We refer to this model as the “direct selection” model because it is could arise from direct selecting acting on the trait (but see below). In the second model (Figure 6b and Supplementary Table 7), there are no large-effect genes and different genes have different levels of constraint. We refer to this model as the “pleiotropic selection” model. We did not consider a model with unconstrained large-effect genes, which would lead to a non-polygenic architecture. Further details of the two models are provided in the Methods section.

We analytically inferred the genetic architecture that arises under each model (Methods). Under both models, common variants have smaller per-allele effect sizes than low-frequency variants (Figure 6c-d), concordant with real traits^10,20^; moreover, both models produced a highly polygenic common-variant architecture (Supplementary Table 7). However, polygenicity differed for the two models at lower allele frequencies (Figure 6e-f). SNP-level flattening (i.e. lower polygenicity at lower allele frequencies) was only observed under the direct selection model; polygenicity was ∼30x lower for *de novo* SNPs than for common SNPs (vs. ∼1.1x under the pleiotropic selection model). Similar our results on real traits (Figure 4), low-frequency polygenicity was ∼4x smaller, and low-frequency heritability was also ∼4x smaller (Figure 6e). The concordance between low-frequency polygenicity enrichment and heritability enrichment is expected. If the selection coefficient of a SNP scales with its squared per-allele effect size, then the effect size bound will be constant, as a function of allele frequency, in units of per-SNP heritability. In units of per-allele effect size, the bound is higher for low-frequency SNPs, so fewer variants reach the bound, leading to lower heritability and proportionally lower polygenicity. However, this concordance is not expected to hold for very rare SNPs, whose effect sizes are limited by biological constraints rather than selection.

For a real trait, the difference in polygenicity between common and *de novo* SNPs could be much larger than the ∼30x difference we observed under the direct-selection model, if there are genes with extremely large effect sizes. It could also be slightly smaller; however, the close correspondence between polygenicity and heritability at allele frequencies above 0.5% suggests that the trend (Figure 6e) will not plateau until much smaller allele frequencies. Thus, we expect that there is a >> 4x difference between the polygenicity of common and low-frequency SNPs for most real traits. We caution that the difference between common vs. low-frequency polygenicity (and heritability) may be a poor proxy for the difference between common vs. *de novo* polygenicity when selection is extremely strong. The effects of flattening may saturate, leading to a small difference in polygenicity between low-frequency and common SNPs (despite the large difference between common and *de novo* SNPs). Such saturation may explain why Number of children, though extremely polygenic, has a small difference in polygenicity between common and low-frequency SNPs (Figure 4).

We analytically computed the heritability explained by a gene (gene-heritability) as a function of its effect size, for SNPs at different allele frequencies. Under the direct selection model, common variant gene-heritability was approximately constant as a function of gene effect size (except for genes with near-zero effect), illustrating that flattening can act at the level of genes (Figure 6g); the effect was weaker for rare SNPs. Under the pleiotropic selection model (which has no SNP-level flattening), there was also no gene-level flattening (Figure 6h). We computed the proportion of heritability explained by the top 10% of genes (ranked by effect size), at different MAF strata. This proportion was strongly frequency dependent under the direct selection model, but not under the pleiotropic selection model (Figure 6i-j).

These results suggest that large-effect disease genes are always constrained, and that one way that this constraint could arise is direct selection acting on the disease itself. However, some forms of pleiotropic selection may produce similar effects as direct selection; for example, in the case of schizophrenia, pleiotropic selection on neurodevelopment broadly may mimic the effects of direct selection on schizophrenia specifically. Therefore, our results do not imply that direct selection is more important than pleiotropic selection, and do not contradict models of selection that are primarily pleiotropic^16,18^.

If flattening occurs at the level of genes, then polygenicity should be increased near strongly constrained genes. We estimated the heritability and polygenicity of SNPs within 50kb of 2,990 loss of function-intolerant genes from ExAC (”ExAC genic SNPs”)^31,32^. These SNPs were more strongly enriched for polygenicity (∼2.9x) than for heritability (∼1.7x) (Supplementary Table 8). Compared with all genic SNPs (± 50kb), ExAC genic SNPs had 1.9x (95% CI: 1.7 *−* 2.0x) larger polygenicity enrichment but only 1.3x (95%CI: 1.3 *−* 1.4x) larger heritability enrichment, implying 0.71x (95%CI: 0.66 *−* 0.76x) smaller average effect sizes (Supplementary Table 8). These estimates suggest that ExAC genes are more likely to be causal but also are more strongly constrained relative to their effect size; they confirm that negative selection at the level of genes affects polygenicity at the level of SNPs.

If the GWAS signal of large-effect genes is constrained, then top GWAS loci should include a mixture of weak perturbations to large-effect, strongly constrained genes (like a “canary in a coal mine”^33^) and strong perturbations to small-effect, weakly constrained genes. In particular, common coding variants, representing strong perturbations, may be less likely to implicate large-effect, strongly constrained genes (consistent with increased polygenicity for coding SNPs; Figure 4). We tested this prediction for 37 fine-mapped IBD GWAS loci^34^, comparing the probability of loss-of-function intolerance (pLI)^31^ between genes harboring coding or noncoding causal risk SNPs. Indeed, 0/8 candidate genes containing fine-mapped coding variants had high pLI (≥ 0.9), compared to 12/29 candidate genes near fine-mapped noncoding variants (rank-sum test p = 0.006 for difference; Supplementary Figure 7 and Supplementary Table 9).

## Discussion

Using a new definition of polygenicity and a new method to estimate it, we compared the polygenicity of 33 complex traits across allele frequencies and functional categories. We determined that low-frequency variants have lower polygenicity than common variants and that biologically important functional categories have higher polygenicity in proportion to their higher heritability. These results demonstrate that negative selection not only constrains common-variant effect sizes but flattens their distribution across the genome, explaining the high polygenicity of complex traits. Furthermore, negative selection constrains the genetic signals of biologically important regions and genes.

A recent study^15^ proposed an “omnigenic model” of complex traits: due to densely connected cellular networks, nearly every gene expressed in a relevant cell type contributes to heritability. As a result, most heritability is explained by “peripheral genes” rather than biologically important “core genes”. On the one hand, our results suggest that the omnigenic model is incomplete, and that negative selection can help explain why core genes explain a limited proportion of trait heritability. On the other hand, our results support a potential distinction between core and peripheral genes (which has been debated^15,35^): among a large set of genes with similarly strong common-variant associations, a small subset may have much larger phenotypic effects, and it may be useful to label these as core genes.

Polygenicity has not precluded GWAS from producing biological insights, and our results provide guidelines to accelerate their progress. First, GWAS follow-up studies should prioritize genes with evidence of constraint, since all large-effect genes are constrained. Loss-of-function variants provide a useful metric of constraint^31^; for GWAS, a maximally informative constraint metric might incorporate noncoding variation^36^, gene expression data^37^, and functional predictions^38^. Second, counter-intuitively, follow-up studies should prioritize associations that do not map to coding regions or large-effect regulatory elements, since SNPs with strongly deleterious effects on their target gene are less likely to implicate strongly pathogenic genes. As our ability to interrogate the gene-regulatory effects of GWAS SNPs improves^38-40^, a potential pitfall would be to prioritize GWAS SNPs with the largest regulatory effects. Third, rare-variant based evidence from exome sequencing studies, even if underpowered, can be used to prioritize GWAS genes. Indeed, exome sequencing studies^41-43^ have been viewed as an attractive complement to GWAS^15^, and our results support this perspective. However, moderately rare coding variants may suffer the same limitations as common regulatory variants.

This study has several limitations. First, we have not quantified the polygenicity of rare or *de novo* variants. An exome-sequencing study of schizophrenia^42^ showed that rare-variant signals are sufficiently polygenic that there was greater power to detect significant pathways than to detect individual genes. Our comparison of common vs. low-frequency variant polygenicity provides a conservative lower bound (∼4x) on the effect of flattening, and our evolutionary modeling suggests a much stronger overall effect (∼30x), although we caution that this number may be slightly smaller or much larger, depending on the distribution of gene effect sizes.Second, our definition of *M*_*a*_ is dependent on the amount of LD between causal SNPs, which has both advantages and disadvantages. On the one hand, it makes *M*_*a*_ identifiable; on the other hand, *M*_*a*_ would be easier to interpret if it only depended on the causal effect size distribution. Third, the nonparametric approach that we used to define and estimate polygenicity may not be optimal for every application. A recent study^11^ fit a parametric model involving a mixture of normal distributions; this approach may provide more accurate estimates of missing heritability as a function of sample size, and it may more accurately predict the performance of risk prediction methods that make similar parametric assumptions^44^. Fourth, S-LD4M can produce biased estimates for depleted functional annotations, albeit in unrealistic settings (Supplementary Figure 1). However, this bias does not affect enriched annotations or low-frequency variants, which are not in strong LD with common variants, and we have avoided reporting polygenicity estimates for depleted functional annotations. Fifth, S-LD4M produces noisy estimates for some annotations and traits, making it necessary to perform meta-analyses across well-powered traits, which may not be representative of all traits. Comparisons of heritability enrichment and polygenicity enrichment are not biased by this filtering process, because the same traits are used to estimate both types of enrichment. Sixth, S-LD4M can potentially be biased due to population stratification, and it should only be applied to data sets where stratification is well controlled. In particular, S-LD4M assumes that any stratification leads to uniform inflation of χ^2^ statistics; this assumption could be violated at loci under positive selection, such as at the LCT locus for height^45^. However, we have only applied S-LD4M to datasets that were corrected for population stratification (including UK Biobank, a relatively homogenous study). Seventh, although our evolutionary modeling supports the hypothesis that flattening affects the distribution of heritability across genes, an alternative explanation is that SNP-level flattening results from increased allelic heterogeneity for common variants near large-effect genes. However, this explanation would require that there be many independent associations per gene. Despite evidence of allelic heterogeneity, it is not a common phenomenon that multiple independent, similarly strong associations implicate the same gene^46^. Eighth, inferences about components of heritability can potentially be biased by failure to account for LD-dependent architectures^17,47-49^. All of our analyses used the baseline-LD model, which includes 6 LD-related annotations^17^. The baseline-LD model is supported by formal model comparisons using likelihood and polygenic prediction methods, as well as analyses using a combined model incorporating alternative approaches^50,51^; however, there can be no guarantee that the baseline-LD model perfectly captures LD-dependent architectures. Despite these limitations, this study advances our understanding of genetic architecture and the evolutionary processes that shape it.

## URLs

Open-source software implementing our method is available at: https://github.com/lukejoconnor/SLD4M. UK Biobank summary statistics are available at: https://data.broadinstitute.org/alkesgroup/UKBB/.

## Acknowledgements

We thank Dr. Benjamin Neale, Dr. Hilary Finucane, Dr. Omer Weissbrod and Dr. Yakir Reshef for helpful comments. This research was funded by NIH grants U01 HG009379, R01 MH101244, R01 MH107649, U01 HG009088 and R01 MH109978.

## Methods

### The effective number of associated SNPs

Here, we provide three definitions of the effective number of associated SNPs (*M*_*a*_). First, we define *M*_*a*_ in terms of the mixed fourth moments of the distribution of causal and marginal effect sizes. This generalizes the definition in Equation (1) (see Overview of methods), which corresponds to the special case of no LD between causal SNPs. Second, we provide an alternative definition (under a random effects model) that formalizes the illustration in Figure 2a, involving the average unit of heritability explained by a causal SNP (see Overview of methods). Third, we provide a concise definition that provides less intuition. We show that these definitions are equivalent in the Supplementary Note.

First, let ***β*** denote the random vector of causal effect sizes for common and low-frequency SNPs. Let *R* be the fixed LD matrix, and let ***α*** = *R**β*** denote the random vector of marginal effect sizes. We use the notation α, *β* to denote randomly chosen entries of ***α***, ***β***. The heritability is *h*^2 =^ E(***β***^*T*^*R****β***) = *M*E(α*β*). *M*_*a*_ is defined as:

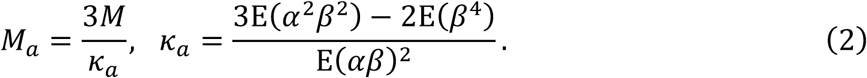

Another possible definition of polygenicity is the effective number of *causal* SNPs, denoted *M*_*b*_:

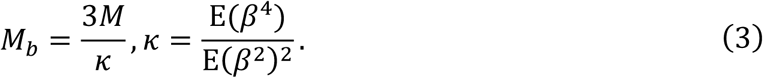

However, this definition is unidentifiable without making a strong assumption (roughly, that linked SNPs have independent causal effect sizes; see below). This definition is equivalent to equation (1) in the Overview of methods section, which defined *M*_*a*_ (rather than *M*_*b*_) in the special case of no LD between causal SNPs; in that case, *M*_*a*_ = *M*_*b*_ (because α = *β* whenever *β* ≠ 0). *M*_*b*_ is different from *M*_*c*_, the number of causal SNPs, except when causal effect sizes follow a point-normal distribution. Because it is implausible that causal effect sizes follow a normal distribution, we view the distinction between *M*_*a*_ and *M*_*c*_ as having greater importance than the distinction between *M*_*a*_ and *M*_*b*_.

Second, we consider a non-i.i.d. normal model:

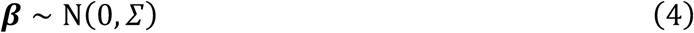

where *Σ* is a fixed diagonal matrix with diagonal entries 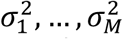. This flexible model generalizes the point-normal model. Because *Σ* is diagonal, 2 is equal to Tr(*Σ*). Note that if *Σ* ∝ *I*, then *κ* is equal to 3 and *M*_*b*_ is equal to *M*. Above, we used the notation E(.) to denote a uniform average across SNPs. We use the notation 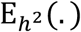 to denote an average across components of heritability, i.e. a weighted average where the probability of choosing SNP *i* is equal to 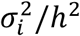. Using this notation, we refer to 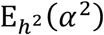as the *average unit of heritability*, and *M*_*a*_ is equal to:

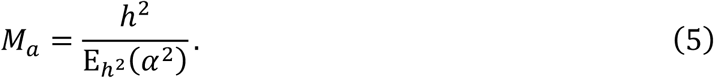

*M*_*b*_ can be defined in a similar manner:

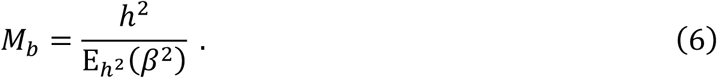

Third, *M*_*a*_ can also be defined without specifying that *Σ* is a diagonal matrix. (The assumption that *Σ* is diagonal is similar to the common assumption that E(***β***^*T*^*R****β***) = E(***β***^*T*^***β***)^2,22^.) Let 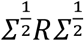. In the diagonal case, S is a weighted LD matrix, where the rows and columns are weighted by the expected effect size of each SNP. The heritability is equal to Tr(*S*) (in the diagonal case, Tr(*S*) = Tr(*Σ*)). *M*_*a*_ is equal to:

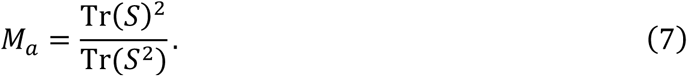

This definition, though concise and natural, provides little intuition.

### Properties of M_a_

*M*_*a*_ has notable mathematical properties.

- **Identifiability**. If there are two SNPs in perfect LD, there is no way to tell based on GWAS data whether one or both of them are causal. If a definition of polygenicity differs depending on whether one or both are causal, then it is unidentifiable. Indeed, *M*_*b*_ (as well as *M*_*c*_) is larger if both SNPs are causal than if only one SNP is causal. However, because both SNPs have identical values of *E*(α^2^), the value of 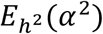 (and therefore *M*_*a*_) is unaffected.
- **Heritability explained by genome-wide significant SNPs**. *M*_*a*_ gives an upper bound on the proportion of heritability explained by SNPs whose effect sizes exceed a specified threshold *T*, such as the genome-wide significance threshold. This proportion can be denoted 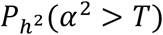. Because α^2^ is nonnegative,

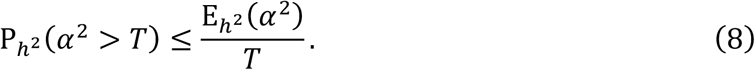

This bound is relatively tight when 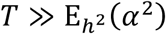. When T is a significance threshold (for example, 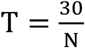 corresponds to genome-wide significance), this bound is most relevant at low sample size.
- **Polygenic prediction accuracy.** Intuitively, increased polygenicity makes prediction more difficult. If *Σ* is given, there is a simple expression for the optimal risk prediction accuracy; this expression provides an upper bound on prediction accuracy in the case that Σ is not given:

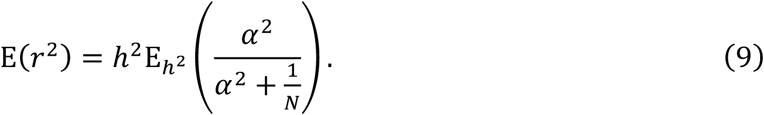

(See Supplementary Note for derivation). At large *N*, prediction accuracy converges to 2. At small *N*, it is approximately a linear function of sample size with slope inversely proportional to *M*_*a*_:

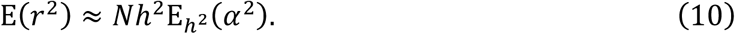

In practice, polygenic prediction is usually performed using large datasets for which this approximation is not appropriate.
- **Effective number of independent SNPs.** Under an infinitesimal model, where every SNP is causal with a normally distributed causal effect size, *M*_*a*_ is equal to the effective number of independent SNPs^21^ (*M*_*e*_). *M*_*e*_ is defined as the number of SNPs divided by the average LD score, or in notation similar to equation (7), as

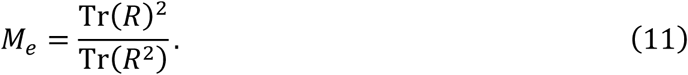

We note that *M*_*a*_ might be close to *M*_*e*_ even when *M*_*b*_ is much smaller than *M*. For example, if the genome comprises perfect LD blocks of 100 SNPs each, then *M*_*a*_ can be equal to *M*_*e*_ even if only 1% of SNPs (1 per LD block) is causal. We also note that the value of *M*_*e*_is strongly dependent on allele frequency, as rare SNPs (which have less LD) contribute strongly to *M*_*e*_. In contrast, *M*_*a*_ will not diverge if many rare SNPs are included, as these SNPs explain little heritability.
- **Symmetry.** *M*_*a*_ is symmetric with respect to the two fixed parameters in the random effects model, *R* and Σ: *M*_*a*_(*R*, Σ) = *M*_*a*_(Σ, *R*). This property is shared by the LD-dependent definition of heritability, E(***β***^T^*R****β***) = Tr(*S*). It is not shared by *M*_*c*_ or by the LD-independent definition of heritability, *E*(***β***^T^***β***).

## Regression equation used in stratified LD fourth moments regression

S-LD4M is justified by an approximate regression equation, which states that the expected value of α^4^ for SNP *i* is approximately proportional to the LD fourth moment of SNP *i*, with a proportionality constant that can be used to estimate *M*_*a*_.

Let *ℓ*^(2)^, *ℓ*^(4)^denote the LD second moment (LD score^28^) and LD fourth moment, respectively, for a randomly chosen SNP:

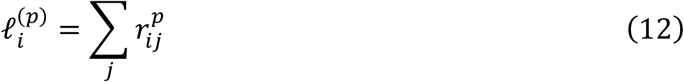

The regression equation is:

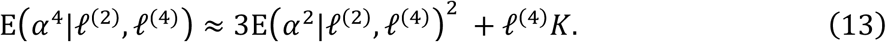

In the first term, E(α^2^*|ℓ*^(2)^, *ℓ*^(4)^) = *ℓ*^(2)^τ, where the coefficient τ is the variance of causal effect sizes^22,28^. In the second term, *K* is related to *M*_*a*_:

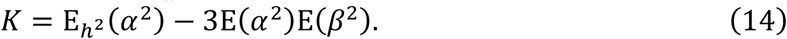

Note that there are three kinds of expectations. First, E(α^2^*|ℓ*^(2)^, *ℓ*^(4)^) and E(α^4^*|ℓ*^(2)^, *ℓ*^(4)^) are conditioned on LD. Second, E(α^2^) and E(β^2^) are not conditioned on LD; rather, they represent unweighted averages over all reference SNPs. Third, 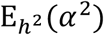 is the average across components of heritability (not uniformly across SNPs).

In the case that there are *P* functional annotations, 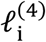 and Κ are vectors of size 1 × *P* and *P* × 1 respectively. Similar to S-LDSC, we make an implicit additivity assumption for the fourth moments of SNPs in the intersection of annotations. (Our simulations violate this assumption, but it does not appear to result in bias).

This regression equation relies on an LD approximation. Roughly, the LD approximation states that if SNP *i* is in LD with causal SNP *j*, then the expected marginal effect size of SNP *i* is approximately proportional to the marginal effect size of *j.* This approximation is exact in important special cases, suggesting that it will be robust in practice. First, we consider the stronger approximation that would be needed to estimate *M*_*b*_. We would need to assume, roughly, that linked SNPs have independent causal effect sizes. More precisely, we would need to assume that the contribution of causal SNP *j* to the marginal effect size of SNP *i*, not conditional on other SNPs, is proportional to the causal effect size of SNP *j*:

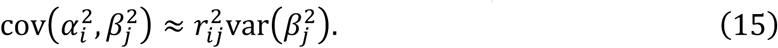

We weaken assumption (15) to obtain our LD approximation:

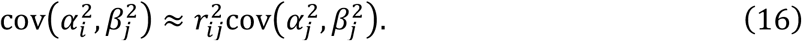

Now, we are assuming that the contribution of causal SNP *j* to the marginal effect size of SNP *i* is proportional to the *marginal* effect size of SNP *j*. Note that (15) implies (16) by applying (15) to the case *i* = *j*. We show that (16) implies (13) in the Supplementary Note. (16) is not violated when other causal SNPs *k* are in strong LD with SNP 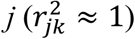, since they would contribute to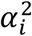in proportion to their contribution to 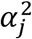 (because 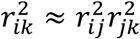). However, it could be violated if there were other causal SNPs in weak LD with SNPs *i* and *j* (violating 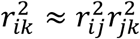).

Equations (15-16) have second-moment analogues that can be used to justify LD score regression^22,28^. LDSC has been justified by assuming that correlated SNPs have uncorrelated effect sizes; this assumption can be stated as:

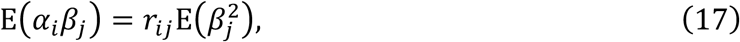

which is analogous to (15). This assumption allows LDSC to estimate a non-LD-dependent definition of heritability, *ME*(*β*^2^). However, a weaker assumption is also possible, analogous to (16):

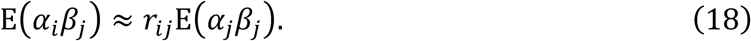

This weaker assumption still holds when perfectly correlated SNPs have correlated causal effect sizes. It allows LDSC to estimate E(α*β*), which is proportional to β^T^*R*β, as follows:

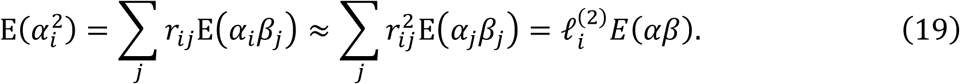

Thus, although LDSC has previously been justified using the assumption that correlated SNPs have uncorrelated effect sizes, only this weaker assumption is strictly necessary in order to estimate an LD-dependent definition of heritability.

## Stratified LD fourth moments regression

In practice we do not observe α, but rather a noisy estimate 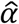. We can correct for sampling noise using:

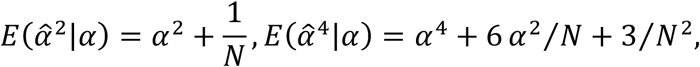

where in practice we use the LD score regression intercept divided by N, denoted 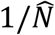, instead of 1/*N*. Let 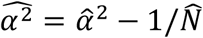 and 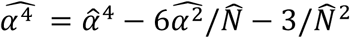.

We perform a two-step inference procedure. First, we use a slightly modified version of S-LDSC^22^ (see below) to estimate 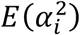 and 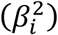 for each SNP, conditional on their respective LD scores and annotation values. Second, we regress the *M* × 1 vector 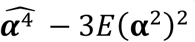on the *M* × *P* matrix ***l***^(4)^ to obtain an estimate of the 1 × *P* vector *K*. The regression is weighted: the weight of SNP *i* is 1 divided by the LD fourth moment of SNP *i* (to all common and low-frequency SNPs). For each annotation *A*_*p*_, we subtract 3*E*(*E*(α^2^)*E*(*β*^2^)*|A*_p_) from *K*_*p*_ to obtain an estimate of *L*, which we combine with the S-LDSC heritability estimate for each category to estimate *M*_*a*_for each category.

Our modified version of S-LDSC uses a slightly modified weighting scheme and does not exclude large-effect SNPs. The regression weight of each SNP is 1 divided by the LD score for that SNP to all common and low-frequency SNPs; this choice is similar to the original version of S-LDSC^22^, but slightly modified for consistency with the weights used in S-LD4M. (We do not exclude large-effect SNPs because these SNPs are important for estimating fourth moments; their exclusion would lead to upwardly biased estimates of *M*_*a*_.)

## Simulations

We simulated summary statistics directly using the asymptotic sampling distribution^23^, rather than by simulating individual-level data. Specifically, the distribution of the summary statistic vector, 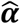, as a function of the true causal effect size vector ***β***, was:

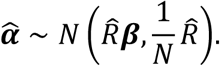

The estimated LD matrix 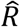 was equal to *I* in simulations with no LD. In simulations with LD, 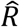 was computed from UK Biobank data (N=460k). Sample correlations were computed for all typed and imputed SNPs (M=1.0M) within 0.1cM of each other on chromosome 1. Blocks of 5,000 SNPs were used, and *R* was set to zero outside of the blocks. These choices were necessary for 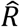 to be stored in memory, due to the large number of SNPs. To ensure that 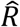 was positive semidefinite, negative eigenvalues were discarded, as previously described^52^.

### Simulations with no LD

We simulated 50,000 total SNPs, 9600 small-effect SNPs, and 400 large-effect SNPs. For small- and large-effect SNPs, effect sizes were drawn from a normal distribution with mean zero and variance 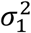and 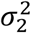respectively; in the three simulations (corresponding to the three example architectures in Figure 2a),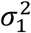was equal to 1/10,000, 1/1,200, and 1/600 respectively.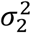was equal to 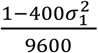, resulting in a heritability of 1. (We note that in all simulations, only *Nh*^2^ affects the results; for example, identical results are obtained at *h*^2^ = 1 and *N* = 20k and at *h*^2^ = 0.2 and *N* = 100k.) We performed simulations atn *N* = 5,000, 25,000, 125,000 and 625,000.

### Simulations with LD

We two sets of simulations with real LD (see above). First, we performed simulations with no functional annotations. We specified that a certain percentage of common SNPs (MAF>0.05) and a certain percentage of low-frequency SNPs (0.05>MAF>0.005) were causal. Conditional on being causal, effect sizes were drawn from a mixture of two normal distributions, with probability *p*_1_ = 0.1 and *p*_2_ = 0.9, and variance 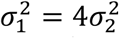. The variance parameter was frequency-dependent; it was proportional to [*p*(1 *− p*)]^0.25^, where *p* is the allele frequency.

Second, we performed simulations with 5 real functional annotations (coding, enhancer, promoter, DHS, repressed). We specified a different probability of being causal for SNPs in each category: 1/2, 1/4, 3/20, 3/80, and 1/20, for SNPs in each annotation and SNPs in no annotation at all, respectively. When a SNP was in multiple annotations, the probability of being causal was the maximum of the respective values. Conditional on being causal, SNPs had independent and identically distributed effect sizes; their effects were drawn from a mixture of two normal distributions, with probability *p*_1_ = 0.1 and *p*_2_ = 0.9, and variance 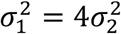. The variances were scaled so that the total (expected) heritability was 0.2.

## Evolutionary modeling

In our evolutionary modeling, each SNP affected one gene and each gene affected both the trait and fitness. We specified an effect size distribution of effect sizes for SNPs on genes (identical for every gene) and a joint distribution for the effect size and selection coefficient of each gene. The effect of a SNP on a trait was its effect on its gene times the gene effect size; the selection coefficient of a SNP was its effect on its gene times the gene selection coefficient. Using an analytical formula for the distribution of allele frequencies conditional on selection coefficients, we obtained the joint distribution of trait effects and allele frequencies for each gene. In detail:

Under each model, we specify a joint distribution of gene effect sizes, denoted *β*_gene_, and selection coefficients, denoted *S*_*gene*_. For the direct selection model, *S*_*gene*_ was equal to 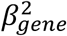, and *β*_gene_ followed a mixture of normal distributions; 95% of genes had variance 10^−3^, and 5% of genes had variance 10^−1^. As a result of this choice, the *de novo* effect size distribution is dominated by large-effect genes. For the pleiotropic selection model, *S*_*gene*_ was independent of *β*_gene_; *β*_gene_ followed a normal distribution with variance 10^−3^ (so that there were no large-effect genes), and *S*_*gene*_ followed a gamma distribution with parameters *k* = 5*/*2 and *θ* = 1*/*1,250.

Under each model, we specify a distribution of effect sizes for SNPs on each gene, denoted *β*_S*NP→*gene_. Each SNP affected one gene, and the distribution was identical for every gene. For the direct selection model, we specified an inverse-gamma distribution with parameters *k* = 100 and *θ* = 1. This is a heavy-tailed distribution; it causes the lines in Figure 6g to plateau, rather than decreasing after attaining a maximum. For the pleiotropic selection model, we specified a mixture of normal distributions; 75% of SNPs had variance 0.2, and 25% of SNPs had variance 0.02. This choice causes the effect size distribution to be highly polygenic, despite the lack of flattening in this model; it also leads to an appropriate relationship between allele frequency and per-SNP heritability.

The effect size of a SNP on the trait, denoted *β*_*SNP*_, was equal to *β*_S*NP→*gene_*β*_gene_. The selection coefficient of a SNP, denoted *S*_S*NP*_, was equal to 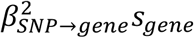. For each gene, we computed the joint distribution of *β*_S*NP*_ and allele frequency *p*, using the formula for the probability density of *p* conditional on *S*_S*NP*_:

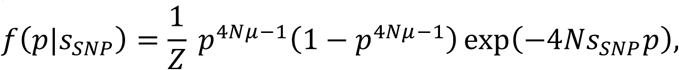

where *Z* is a constant, *N* = 100, and *μ* = 1*/*200 (so that the exponent 4*Nμ −* 1 is equal to 1). We note that the model is overparameterized; for example, identical results would be obtained with 100x larger *N* and 100x smaller *S*_*gene*_ and *μ*.

We simulated 20,000 genes. For each gene, we computed the second and fourth moments of *β*_*SNP*_ for SNPs at allele frequencies *p* = 0.25, 0.05, 0.01,0.002,0.004, by multiplying the probability density function of *β*_*SNP*_ by *f*(*p|S*_*SNP*_) and approximating the integral using a sum over 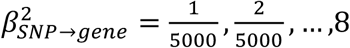. Averaging over genes, we computed the variance in per-allele effect sizes (Figure c-d), the per-SNP heritability (variance times heterozygosity; Figure 6e-f), and the polygenicity (inverse of kurtosis; Figure 6 e-f). We also computed the proportion of heritability explained by the 10% of genes with largest 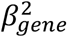(Figure 6i-j).

We also computed the heritability explained by a gene as a function of 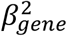(Figure 6g-h). In Figure 6g, *S*_*gene*_ was equal to 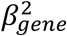; in Figure 6h, *S*_*gene*_ was fixed at 2 × 10^−4^. The heritability explained by the gene at a given allele frequency was equal to the variance of 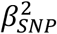 at that allele frequency, times the heterozygosity.

---

**Supplementary Figure 1.**
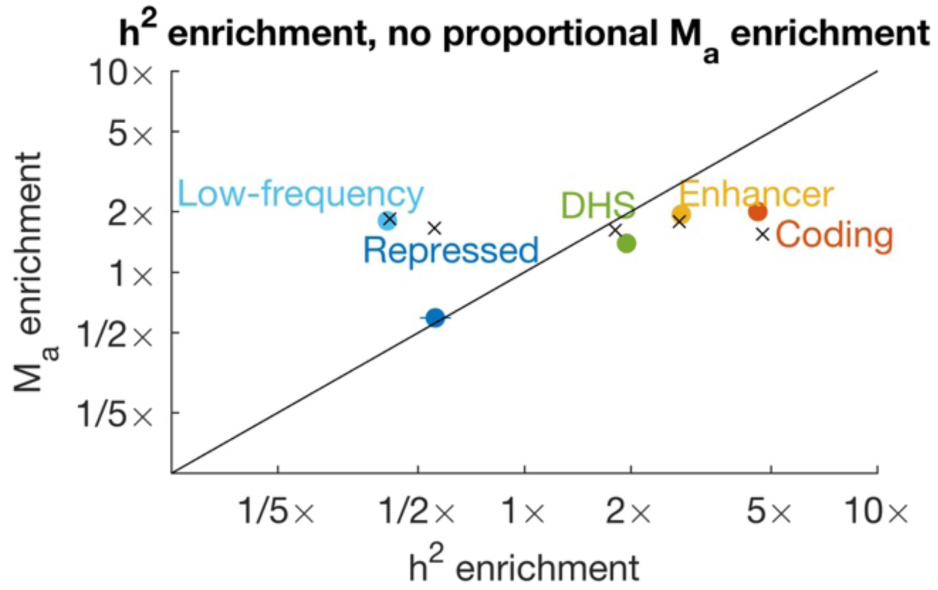
Simulations with heritability enrichment and no proportional polygenicity enrichment. Due to some sparsity being driven by differences in heritability enrichment across categories, there is lower sparsity (polygenicity enrichment) within each individual category than within their union. We observed strong downward bias for the repressed category, which is depleted for heritability; we hypothesize that this bias is the result of imperfect resolution to distinguish LD to this category from LD to nearby SNPs, which leads to inflated estimated fourth moments because the nearby SNPs have larger causal effect sizes. This bias has little effect on our estimates for categories depleted for heritability because the nearby SNPs have smaller causal effect sizes than the SNPs in the category, and therefore very little effect on fourth moments; it also has little effect on our estimates for low-frequency SNPs because these SNPs are never in strong LD with common SNPs. In analyses of real traits, we do not report polygenicity enrichment estimates for categories that are depleted for heritability. Based on 1,000 simulations. Error bars indicate 95% confidence intervals. Numerical results are contained in Supplementary Table 1.

**Supplementary Figure 2.**
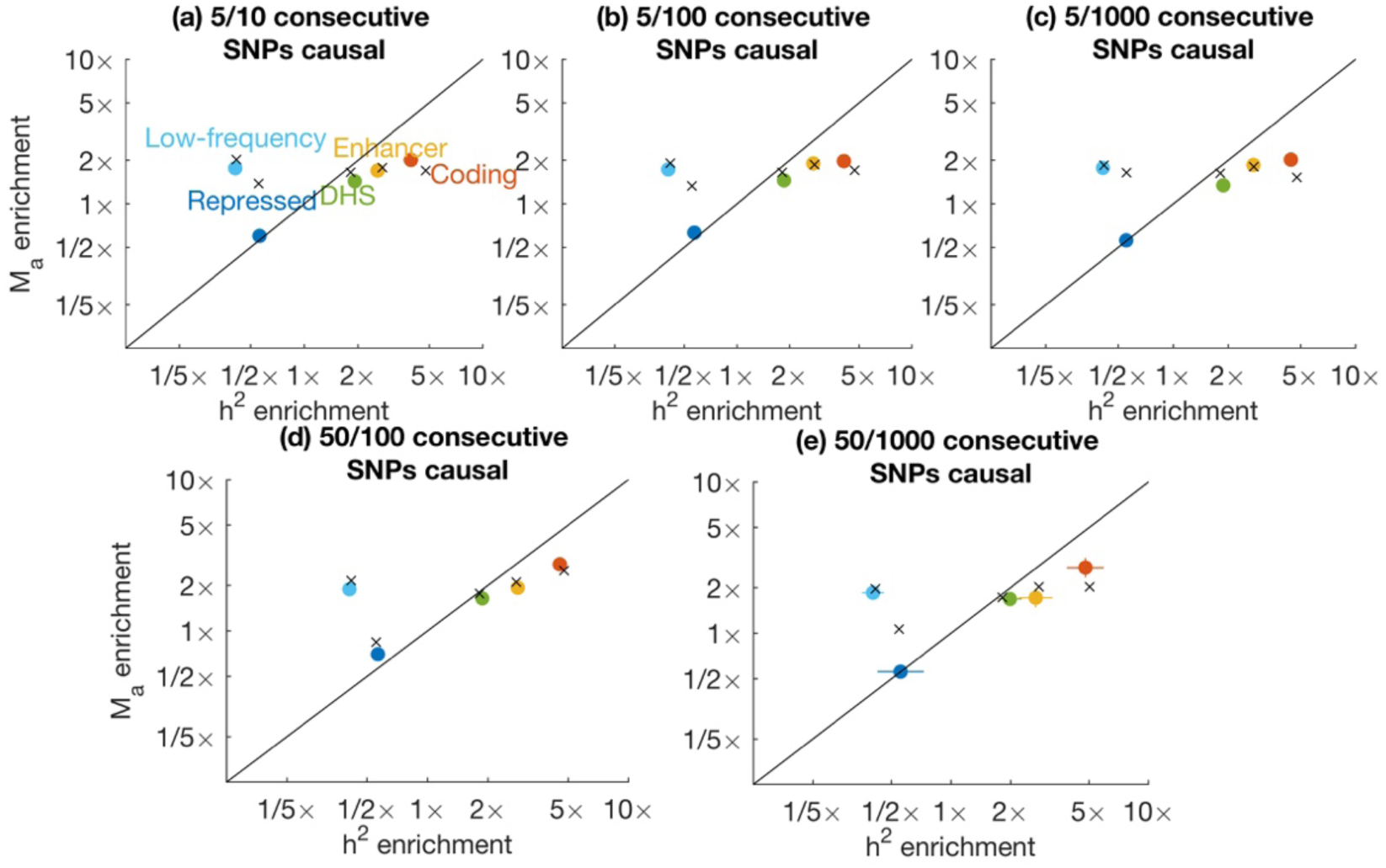
Simulations with clustering of causal SNPs. The probability for a SNP to be causal was zero for most of the genome, and nonzero for contiguous blocks of SNPs of different sizes. (a) Blocks of 10 SNPs with a 50% chance of being causal. (b) Blocks of 100 SNPs with a 5% chance of being causal. (c) Blocks of 1,000 SNPs with a 0.5% chance of being causal. (d) Blocks of 100 SNPs with a 25% chance of being causal. Based on 1,000 simulations. Error bars indicate 95% confidence intervals. Numerical results are contained in Supplementary Table 1.

**Supplementary Figure 3.**
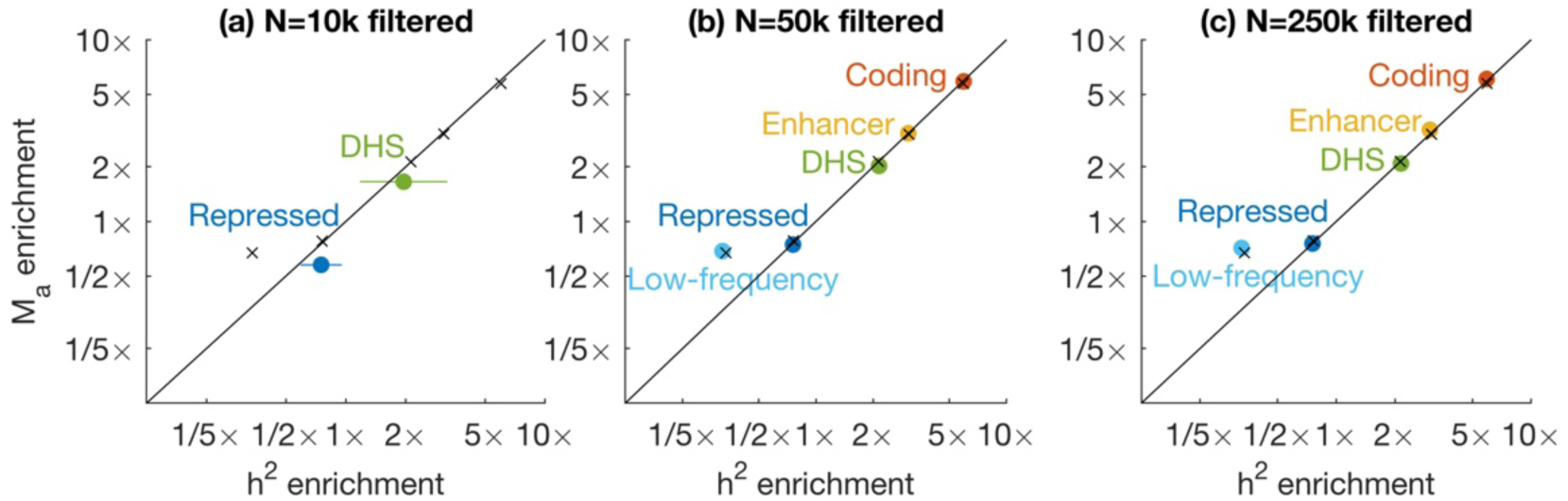
Simulations with ascertainment based on power. For each annotation, simulation runs were discarded if heritability for that annotation was not significantly different from zero (*Z* > 2) or if the standard error of the *M*_*a*_ estimate for the annotation was larger than two times the median *M*_*a*_ point estimate. Results are not shown for annotations where the fraction of retained simulations was less than 1%, and confidence intervals are large for annotations where the fraction of retained simulations is low. See Supplementary Table 2 for the fraction of simulations that were retained in each case. Based on 1,000 simulations (before filtering). Error bars indicate 95% confidence intervals. Numerical results are contained in Supplementary Table 1.

**Supplementary Figure 4.**
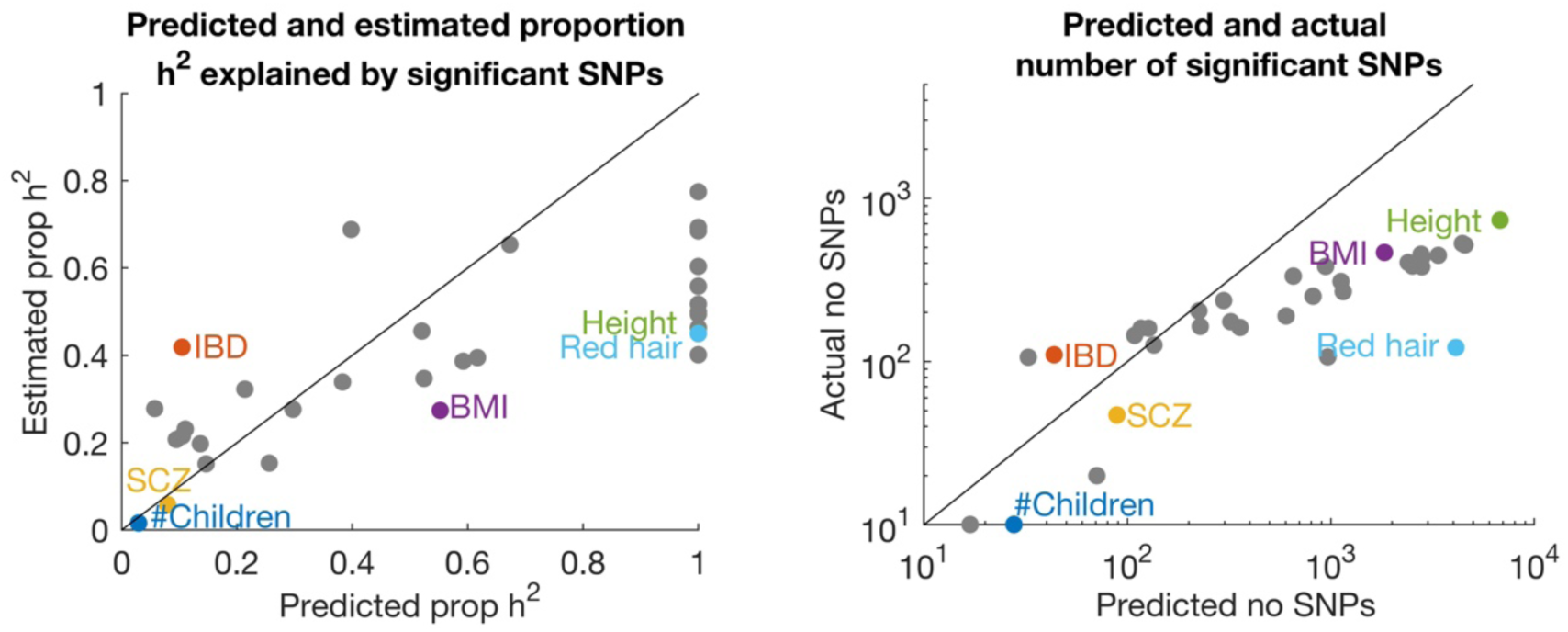
Heritability explained by genome-wide significant SNPs. *M*_*a*_ gives a predicted upper bound on the proportion of heritability explained by significant SNPs; similarly, it also gives an upper bound on the number of significant SNPs (Methods). Significant SNPs (χ^2^ > 30) were chosen using a greedy pruning procedure, with a gap of at least 0.5cM between them. We estimated the proportion of heritability explained by these SNPs as the sum of their estimated marginal effect size magnitudes. We caution that this estimate may be upwardly biased due to winner’s curse and due to subtle LD between these SNPs.

**Supplementary Figure 5.**
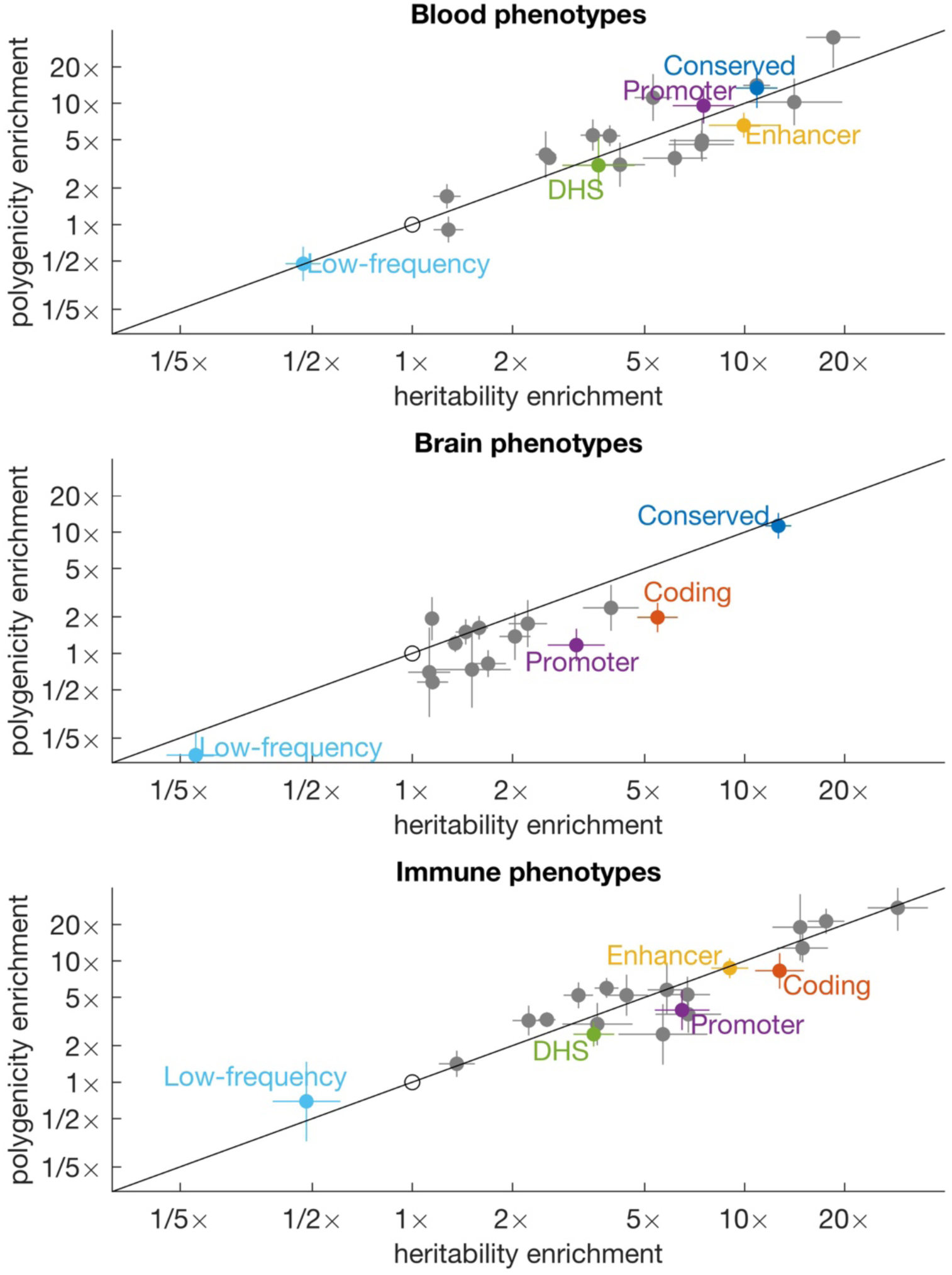
Heritability and polygenicity enrichment meta-analyzed across related traits. There were 6 blood-related phenotypes (eosinophil count, platelet count, platelet distribution width, RBC count, RBS distribution width, and WBC count); 8 brain-related phenotypes (Alzheimer’s, BMI, college, morning person, neuroticism, smoking status, number of children and schizophrenia); and 5 immune-related phenotypes (all autoimmune, asthma, eczema, IBD, and RA). Results for each annotation are meta-analyzed across well-powered traits within each group, and each annotation is plotted if at least three traits had a well-powered polygenicity estimates for that annotation.

**Supplementary Figure 6.**
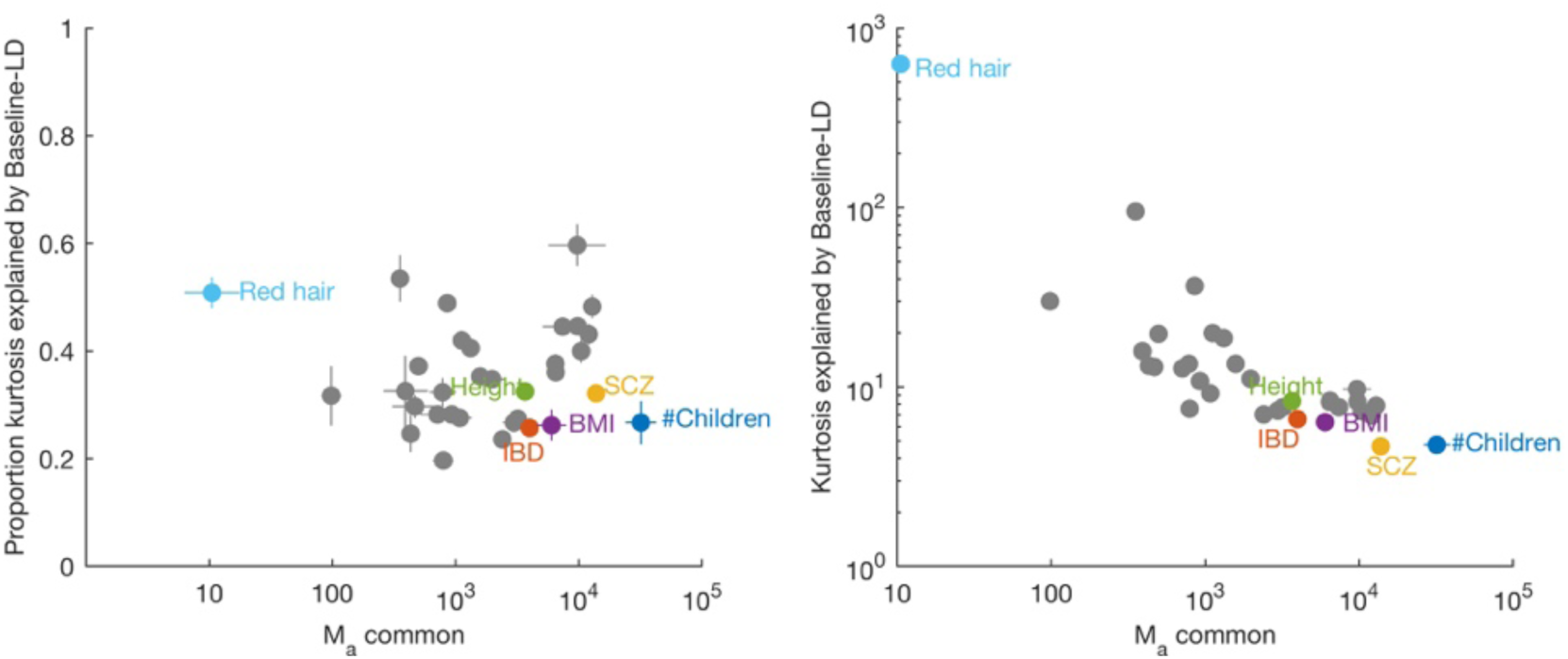
Common-variant sparsity explained by baseline-LD model. Sparsity is defined as *M*_*e*_*/M*_*a*_. Some sparsity is expected due to differences in per-SNP heritability across categories; S-LDSC was used to estimate the causal effect size of each regression SNP, and these estimates were multiplied by the observed χ^2^ statistics to estimate the sparsity explained by the model. (a) The proportion of kurtosis explained was defined as the ratio of logs (log-sparisty explained divided by log-sparsity). It ranged between 20% and 50% for most traits, and there was no clear tendency for traits with greater polygenicity to have greater or smaller percent sparsity explained. (b) The amount of sparsity explained was strongly negatively correlated with polygenicity (and positively correlated with total sparsity).

**Supplementary Figure 7.**
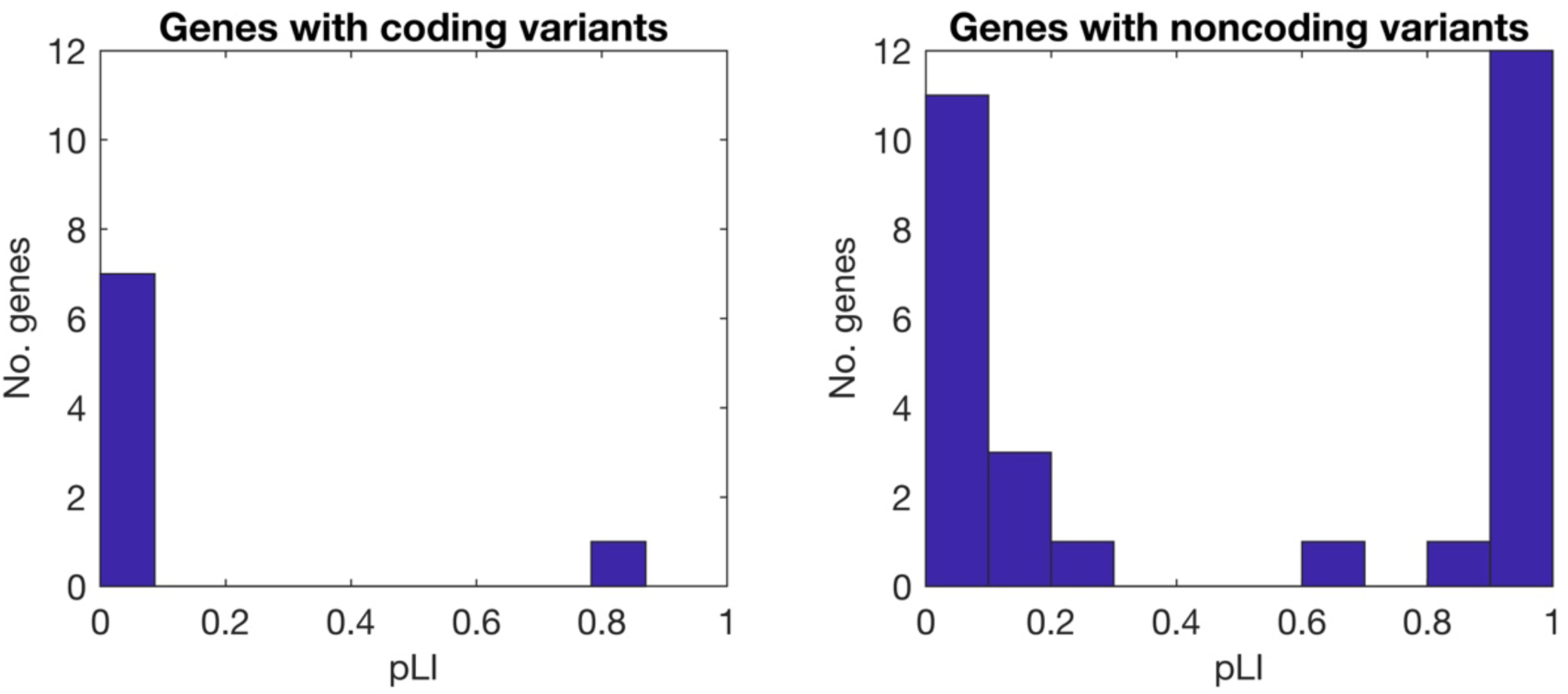
Distribution of pLI values for IBD genes with fine-mapped coding and noncoding variants. Causal variants for Crohn’s disease and ulcerative colitis were fine mapped in ref. ^34^. 47 fine-mapped variants with >50% posterior probability and 1-2 annotated protein-coding genes were selected. There were 7 SNPs (all noncoding) with 2 annotated genes, and pLI values were similar (within 0.02) for 6/7 genes; pLI values were averaged for these loci. There were 3 genes with multiple causal variants (ranging from 2-6), including both coding and noncoding variants; these were counted only once, as coding genes. pLI values were significantly smaller for genes with coding variants than for genes with noncoding variants (single-tailed rank-sum test *p* = 0.006). We caution that pLI values are computed based on allele frequencies of loss of function variants, although high pLI genes may have common missense variants. SNP-by-SNP data is contained in Supplementary Table 7.

**Supplementary Table 1.**
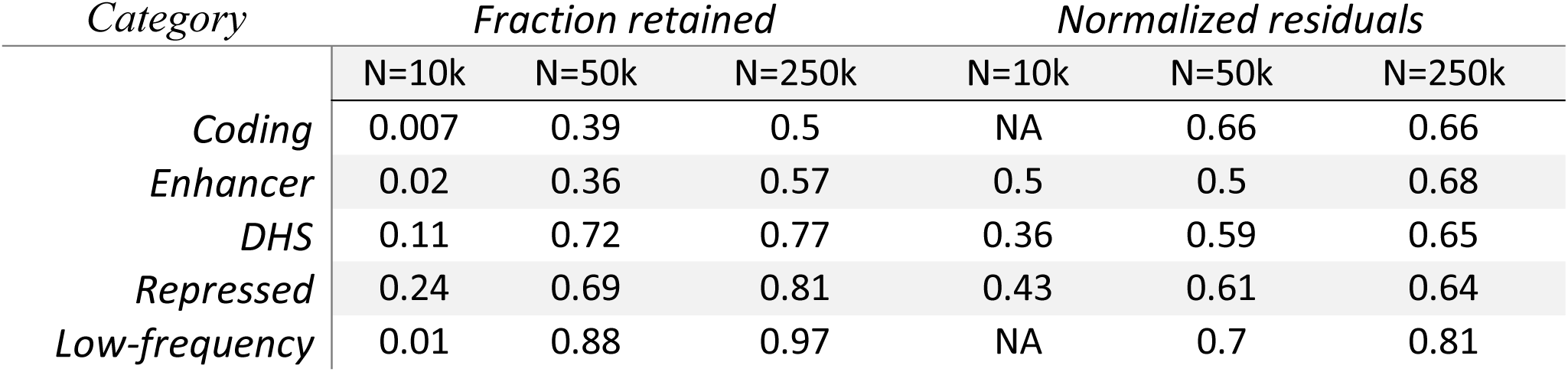
(See Excel file) Numerical results of simulations. Sheets (a)-(f) correspond to Figure 1a-c and Supplementary Figures 1-3, respectively.

**Supplementary Table 2.** Ascertainment and standard errors in simulations at different sample sizes. For each annotation, simulation runs were discarded if heritability for that annotation was not significantly different from zero (*Z <* 2) or if the standard error of the *M*_*a*_ estimate for the annotation was larger than two times the median *M*_*a*_ point estimate. For each annotation, the fraction of simulation runs that were ascertained is reported, as well as the standard deviation of the normalized residuals for the ascertained simulations. The normalized residuals are defined as 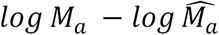 divided by the jackknife standard error, and they should be equal to one if the jackknife standard errors are perfectly calibrated. Values less than 1 indicate conservative standard errors. Jackknife standard errors for polygenicity enrichment are nearly proportional to standard errors for Ma, since there is relatively little noise in the denominator (the estimate of *M*_*a*_ for all common and LF SNPs).

**Supplementary Table 3.** (see Excel file) Datasets analyzed. 29 UK Biobank traits were selected to have low pairwise genetic correlations and high power, as measured by the significance of the S-LDSC heritability estimate; 28 of these were analyzed in ref.^20^, and red hair pigmentation was added. Four additional diseases were selected on the basis of availability of summary statistics for low-frequency SNPs. Seven additional datasets were used for replication.

**Supplementary Table 4.** Comparison of common-SNP *M*_*a*_ estimates for traits with multiple available datasets. Standard errors are also reported. No *M*_*a*_ estimates were significantly different (*p <* 0.05 assuming independent errors) for any pair of datasets.

| *Trait* | *Reference* | *N* | *M* | *Common $\log_{10} M_a$* |
| --- | --- | --- | --- | --- |
| *Height* | UKBB | 458k | 10M | **3.56(0.02)** |
| *Height* | Lango Allen et al. 2010 | 131k | 1.0M | **3.49(0.11)** |
| *BMI* | UKBB | 458k | 10M | **3.78(0.12)** |
| *BMI* | Speloites et al. 2010 | 122k | 1.0M | **3.51(0.33)** |
| *College* | UKBB | 455k | 10M | **4.08(0.05)** |
| *Years of education* | Rietveld et al. 2013 | 127k | 1.0M | **4.26(0.18)** |
| *Years of education* | Okbay et al. 2016 | 329k | 1.0M | **4.23(0.09)** |
| *Type II Diabetes* | UKBB | 459k | 10M | **2.85(0.14)** |
| *Type II Diabetes* | Morris et al. 2012 | 61k | 1.0M | **2.85(0.38)** |
| *Neuroticism* | UKBB | 372k | 10M | **3.99(0.23)** |
| *Neuroticism* | Okbay et al. 2016 | 171k | 1.0M | **4.26(0.10)** |
| *Rheumatoid arthritis* | Okada et al. 2014 | 38k | 4.6M | **3.03(0.10)** |
| *Rheumatoid arthritis* | UKBB | 459k | 10M | **3.08(0.20)** |

**Supplementary Table 5.**
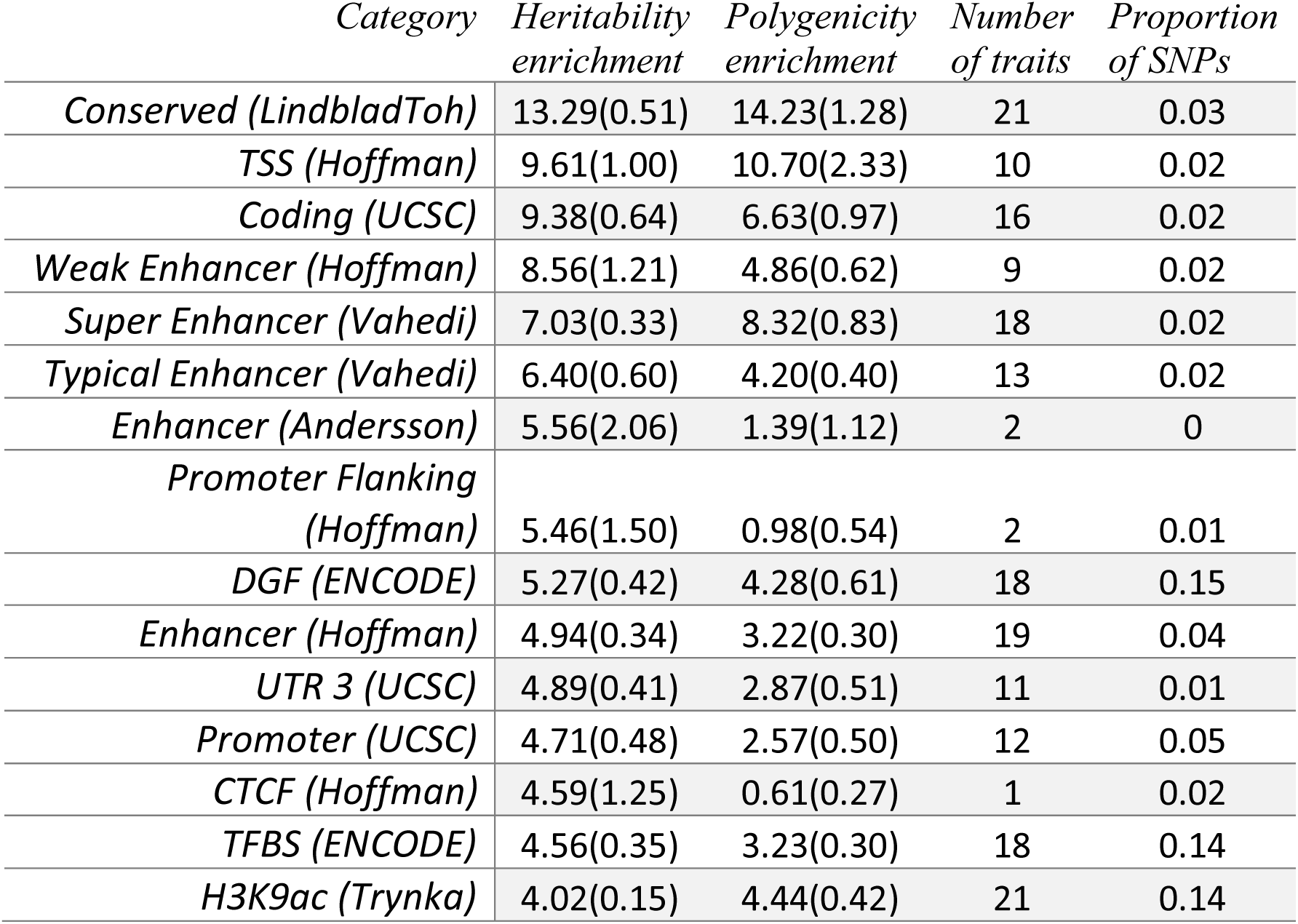
(See Excel file) Complete results of S-LD4M and S-LDSC on 33 traits.

**Supplementary Table 6.**
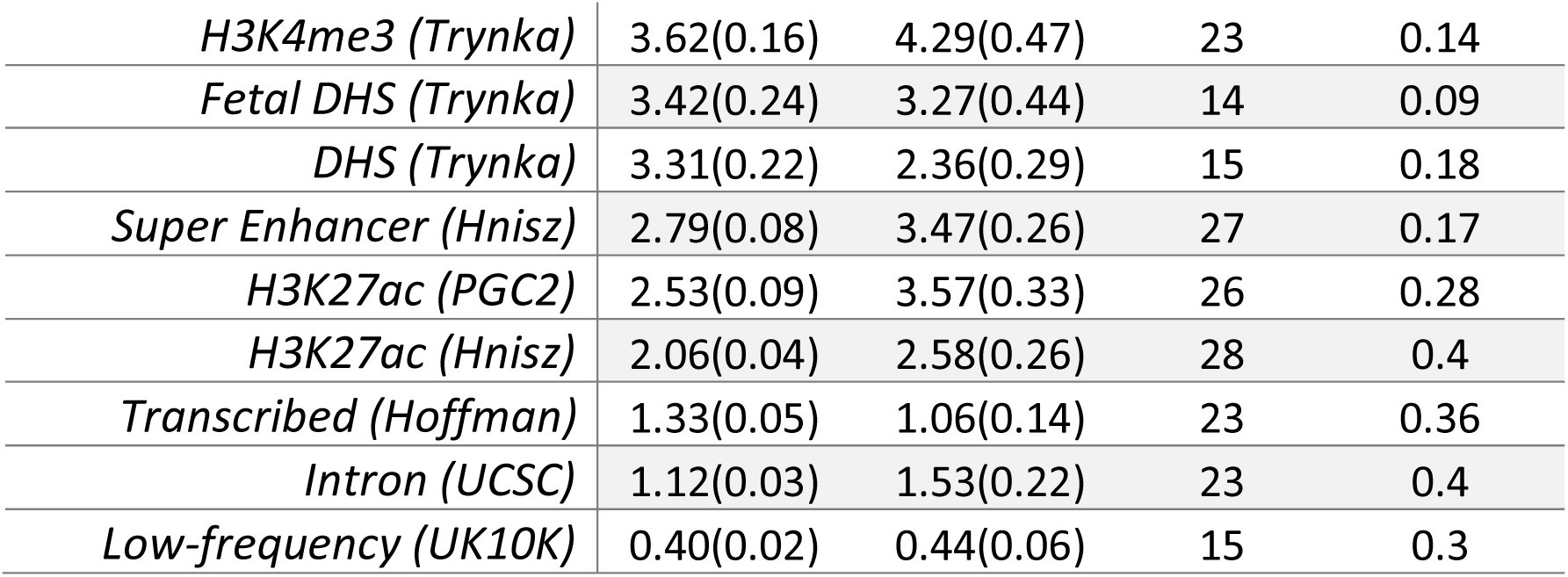
Polygenicity and heritability enrichment for functional annotations, meta-analyzed across well-powered traits. The number of traits used in the meta-analysis is indicated; traits were excluded if the heritability estimate for a trait-annotation pair was not significantly different from zero, or if the standard error on the *M*_*a*_ estimate was greater than 4 times the median point estimate for that annotation across traits. Annotations were excluded if the number of remaining traits was less than 10 or if the meta-analyzed heritability enrichment estimate was less than 1 (except for the low-frequency category). Standard errors are also reported.

**Supplementary Table 7.** (see Excel file) Numerical results from Figure 6.

|  | *Enrichment* | *log<sub>10</sub>Enrichment* |
| --- | --- | --- |
| *Polygenicity* | 1.82 | 0.27 (0.02) |
| *Heritability* | 1.35 | 0.13(0.007) |
| *Average effect size* | 0.71 | -0.15 (0.02) |

**Supplementary Table 8 Polygenicity and heritability enrichment of ExAC genes.** Enrichments are reported for SNPs in and near ExAC LoF-intolerant genes^32^ compared with SNPs near any gene (in and near is defined as the gene body plus or minus 50kb). The average effect size is equal to the heritability divided by the polygenicity (Figure 2; Methods). Standard errors are also reported.

**Supplementary Table 9 see Excel file) Fine-mapped IBD genes from ref.^35^ harboring coding and noncoding variants.** Coding and noncoding SNPs with > 50% posterior probability and 1-2 nearby genes are listed. pLI values are averaged for SNPs with 2 genes. Genes with both coding and noncoding variants are included in Supplementary Figure 7a and not in panel b.

